# The shape of attention: How cognitive goals sculpt cortical representation of speech

**DOI:** 10.1101/2025.05.22.655464

**Authors:** Moïra-Phoebé Huet, Mounya Elhilali

**Affiliations:** Department of Electrical and Computer Engineering, Johns Hopkins University, Baltimore, MD, USA

## Abstract

Perception requires more than passive sensing—it involves prioritizing the features most relevant to ongoing cognitive goals, a process guided by selective attention. A central question is whether attention operates by enhancing all features of a selected target, or by optimizing neural encoding around the specific demands of the task—i.e., is selective attention fundamentally anchored around task targets or around task goals? Here, we recorded electroencephalography (EEG) while participants performed two speech tasks—comprehension and detection—on identical auditory stimuli. Task difficulty was manipulated by introducing controlled background noise that increased cognitive demands without reducing speech intelligibility. We developed a novel EEG-based method, the Modulation Response Function (MRF), which captures cortical sensitivity to spectro-temporal features via spectrogram reconstruction. Behaviorally, comprehension performance declined with increased difficulty, with greater reliance on semantic cues, while detection performance remained near ceiling. Neurally, both envelope tracking and MRF magnitude were higher during comprehension, reflecting greater cognitive engagement. Critically, spectro-temporal tuning differed across tasks: formant-related modulations were selectively enhanced during comprehension, whereas pitch-related modulations were emphasized during detection. These findings support a discriminative model of attention, where cortical encoding is flexibly reshaped according to cognitive goals, selectively amplifying the features most relevant for successful task performance.

## Introduction

### Background

Navigating the sensory world requires more than passively receiving information—it demands an active prioritization of what matters most. Attention is the brain’s solution to this challenge: a flexible mechanism that filters sensory input to meet the brain’s needs. This selective filtering is widely considered a core function of attention, enabling the brain to enhance relevant information while suppressing distractions [1]. Whether we are picking out a familiar face in a crowd or following a single voice in a noisy room, attention allows us to focus, adapt, and extract meaning. These examples highlight that selection is not an end in itself—it is a means to different cognitive ends, from mere detection to rich interpretation. Attention may thus be best seen as a mechanism that aligns perceptual processing with cognitive goals.

This perspective has played a central role in cognitive science, where attention is commonly conceptualized as a filtering mechanism that prioritizes information according to its relevance. Foundational models [2,3] proposed that attention functions by selecting inputs aligned with current goals for further processing, while discarding irrelevant information early in the perceptual stream. While these models underscored the significance of relevance, they left unresolved whether attention uniformly enhance all relevant features, or can it selectively target specific dimensions of a stimulus depending on task demands? This distinction forms the basis for contrasting theoretical perspectives on the mechanisms regarding the mechanisms through which attention shapes sensory encoding.

One influential view in the attention literature proposes that attention acts as a global gain mechanism, enhancing neural responses to all features of a selected sensory object. This perspective is supported by object-based accounts of attention [4]. For instance, in visual attention, attending to a particular spatial location or object typically results in the simultaneous enhancement of multiple stimulus attributes at that location—even when only one attribute is behaviorally relevant [5,6]. A similar effect has been reported in auditory perception: directing attention to one acoustic feature (e.g., spatial location or pitch) appears to enhance the neural representation of all associated features of that auditory object [7,8]. Together, these findings support the notion that attention operates as a broad enhancement mechanism, amplifying all features of a selected object, irrespective of the specific cognitive goal.

However, experimental findings have challenged this uniform gain perspective, suggesting a more flexible, dynamic filtering process that selectively enhances task-relevant sensory features. This view has been primarily supported by research in the visual domain, where studies suggest that attentional modulation may adapt to the specific demands of the task rather than uniformly boosting all features of a selected object or location. For instance, computational and behavioral work has shown that attention can optimize performance not by enhancing the target feature itself, but by amplifying the most informative dimensions—even when these are not aligned with the target [9,10]. Neurophysiological studies further demonstrate that the same visual input can elicit distinct patterns of cortical activation when the task requires detection, identification, or discrimination [11,12]. Similar findings have also emerged in audiovisual speech processing, where the same stimuli evoke distinct neural activity patterns depending on whether participants attend to phonological, semantic, or visual features [13]. Together, these results suggest that attention may act as a dynamic, goal-sensitive filter rather than a fixed amplification mechanism.

From an implementation standpoint, these insights raise a critical design question: What is the objective function that the biological (or a computational) system aims to deploy in order to reflect the flexible, goal-dependent nature of attention? Computationally, one might envision an objective function that selectively boosts the representation of a specific target class— effectively simulating attentional selection as a pre-processing step that prioritizes certain stimuli for downstream tasks. This would align with the object-based enhancement view, where attention operates as a mechanism to enhance all features of a selected input. However, if attention is better understood as a task-contingent filter that dynamically adapts to behavioral goals, then the objective function may instead need to be structured around task performance itself—optimizing for detection, identification, or discrimination, regardless of the specific target class being attended to. In this case, a system must learn to modulate representational gains based on task context, selectively enhancing the most informative features as needed. This approach shifts the focus from static category selection to dynamic optimization, where attention emerges as a byproduct of performance-driven constraints. The appropriate formulation thus depends on the conceptual framing of attention: is it a mechanism for prioritizing specific content, or for aligning processing with cognitive demands?

Speech provides a particularly compelling framework to reconcile this tension and investigate how attention adapts to different cognitive goals. It naturally supports a wide range of listening objectives—such as detecting whether someone is speaking, identifying the speaker, or understanding the message—making it ideally suited to examine how attentional filtering may shift depending on the listener’s intent. In the auditory domain, it is widely accepted that selective attention enhances the target signal while suppressing background noise [14–16]. This attentional modulation enables a broad range of speech-related operations, among which two stand out for their contrasting cognitive demands: speech detection, which allows us to recognize the presence of a vocal signal without necessarily extracting its linguistic content, and speech comprehension, which engages more elaborate mechanisms to integrate phonetic, lexical, semantic, and contextual information.

While both processes rely on speech perception, they differ in their cognitive demands and likely in how attention contributes to each. When the goal is to detect the presence of a sound, attention improves the detection of target sounds by lowering their perceptual threshold—either by enhancing sensitivity to a specific frequency range [17–19] or to a particular spatial location [19]. At the neural level, this corresponds to sharpened cortical responses to specific acoustic features of the attended sound, a sensory gain mechanism observed in both animals [20] and humans [21,22]. However, in more complex tasks such as speech comprehension, listeners must go beyond mere detection to extract linguistic meaning. Selective attention enables them to focus on a single speech stream amid competing sounds—a phenomenon known as the “cocktail party effect” [23]. This process leverages temporal and spatial cues [24,25], and is reflected neurally in the brain’s ability to encode the attended speech stream. This neural representation can be reconstructed from brain responses using models such as spectro- temporal receptive fields (STRF) and temporal response functions (TRF) [26–28], an approach commonly referred to as neural tracking [29]. More specifically, attention enhances speech intelligibility by modulating cortical activity to track the temporal structure of speech [26] and by refining spectral selectivity to emphasize relevant acoustic features [27]. These findings underscore the importance of spectro-temporal processing in speech processing, highlighting its central role in how speech is represented and understood.

A key framework for understanding spectro-temporal processing in speech perception is the concept of spectro- temporal modulations, often described in terms of modulation power spectrum (MPS) or modulation profile. The MPS describes how the energy of a sound signal varies simultaneously across time and frequency, highlighting the acoustic structures essential for speech [30,31]. As illustrated in Fig 1 the MPS is obtained through a series of transformations. First, a spectrogram is generated from the acoustic waveform representing the sound’s frequency content over time. Then, the spectro-temporal modulations are extracted at each time point, while still integrating information across time to capture temporal modulations. Finally, by averaging these modulations over time, the overall MPS is computed, revealing the key spectral and temporal modulations that contribute to intelligibility.

**Fig 1.**
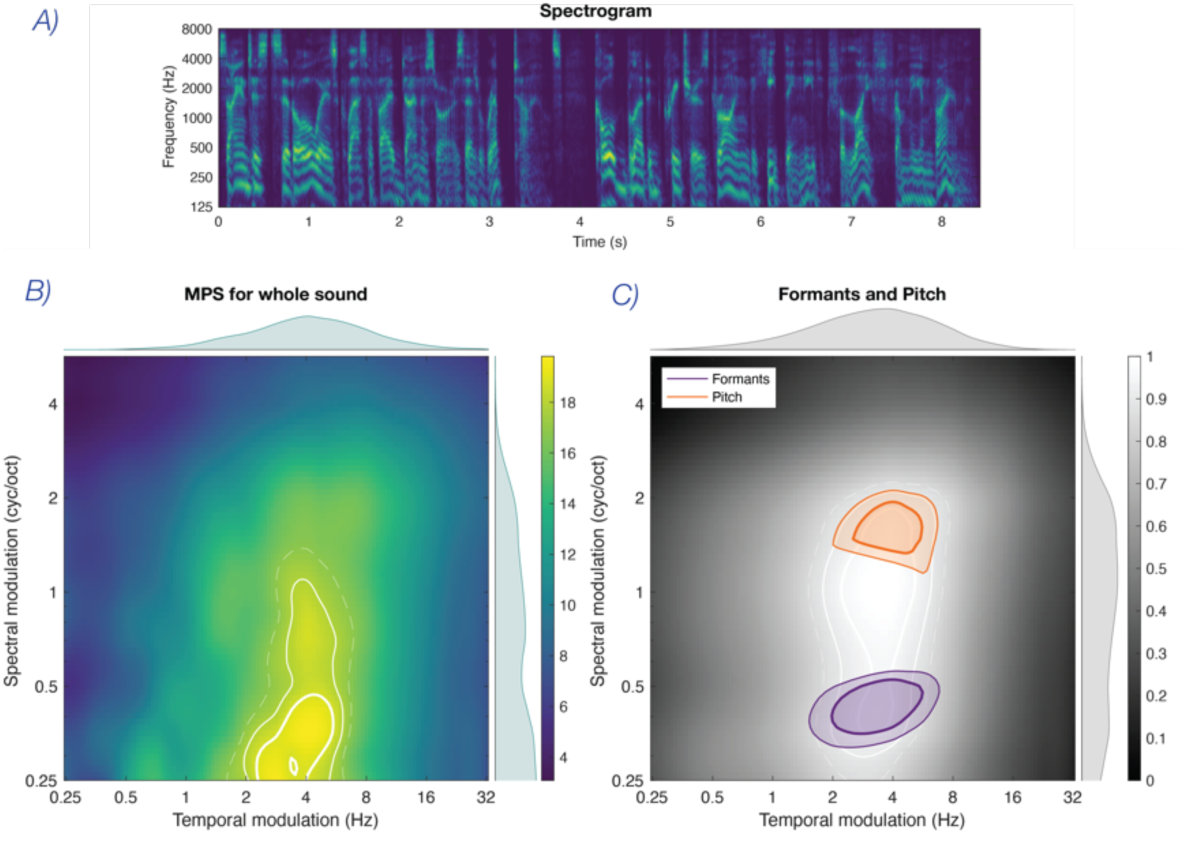
Speech MPS Characteristics. From a spectrogram (Panel A), a modulation profile is derived for each time point t and then averaged across the entire sound (Panel B). Each modulation profile is represented with temporal modulation (in hertz) on the x-axis and spectral modulation (in cycles per octave) on the y-axis. The bold white line indicates the 95th percentile, the solid white line indicates the 90th percentile, and the dashed white line indicates the 85th percentile. The average distribution along the spectral modulation axis is displayed to the right of each profile, while the average distribution along the temporal modulation axis is shown above each profile. Panel C represents the modulation profile of speech from a male speaker (in white) and highlights the regions of the profile corresponding to formants (in purple) and pitch (in orange). While these features align with previous studies [32,33], the exact values may differ due to methodological variations.

Studies have demonstrated that speech intelligibility is commensurate with the preservation of spectro-temporal modulations within specific frequency and temporal ranges [32,34]. Moreover, formants, which are critical for intelligibility, appear in the MPS at low spectral modulations and intermediate temporal modulations, while gender cues—mainly pitch—are found at higher spectral modulations and lower temporal modulations [32,33]. Fig 1 illustrates the regions of the MPS where formants and pitch cues are represented, based on our analysis of male and female voices from approximately 30 hours of audiobook recordings. This comparison allowed us to identify the spectral regions corresponding to formants and pitch cues (see Supplementary Material for details). Importantly, these features do not contribute equally to different speech-related tasks. Pitch-related information—corresponding to the perceptual correlate of the talker’s fundamental frequency (F0)—is particularly relevant for identifying the presence and source of a voice—making it essential for speech detection [35]. In contrast, formant-related modulations are key to phonemic recognition [36] and speech intelligibility [37]—two interdependent processes that support successful speech comprehension. Thus, attention may selectively enhance different parts of the spectro-temporal space depending on the cognitive goal—detection or comprehension.

### Hypotheses

This study examines whether attention operates as a flexible filter that reshapes sensory encoding to align with distinct cognitive goals. While it is well established that attention modulates sensory encoding, studies differ on how this modulation plays out in practice. Neural responses are known to depend on both cognitive demands and sensory features [38], making it essential to isolate the influence of task goals from changes in the stimulus itself. A common limitation across studies is that changing the task often means changing the stimulus too—for example, by altering distractor configurations, perceptual features, or task-relevant targets [9–11]. This makes it difficult to determine whether observed differences in brain activity reflect task-driven attentional filtering or merely differences in the input. This issue is further compounded by the fact that many studies rely on constrained or simplified paradigms. Some use short, low-level stimuli such as tones or syllables [39–41] while others compare tasks that differ in the level of linguistic analysis—such as phonological versus semantic processing [13] or degrees of listening effort [42]. However, these tasks typically fall within the same overarching domain: they all involve processing with the linguistic content of speech. As such, they do not fully capture how attention might operate across qualitatively different cognitive goals—such as distinguishing between simply detecting the presence of speech and understanding its meaning.

Building on these observations, the present study isolates the role of the cognitive goal—detection versus comprehension—by keeping the auditory stimulus strictly constant and varying only the task instruction. This design allows us to test whether attention modulates speech representations in a goal-sensitive manner. Here, we propose two competing hypotheses regarding how selective attention modulates cortical representations of speech during detection and comprehension tasks. The *Gain hypothesis* predicts that attention enhances all spectro-temporal features of speech in a uniform manner, leading to similar cortical responses across tasks, regardless of potential differences in overall magnitude driven by task difficulty or cognitive engagement. In contrast, the *Discriminative hypothesis* suggests that selective attention modulates different spectro- temporal features depending on the cognitive demands of the task—for example, enhancing formant-related features during comprehension and pitch-related features during detection.

To empirically test these hypotheses, we need a method capable of capturing how selective attention modulates neural representations of speech in a task-dependent manner. Previous fMRI studies have reconstructed cortical representations of spectro-temporal modulations from neural responses to speech and other natural sounds [33,34,43,44], but their limited temporal resolution prevents them from isolating rapid acoustic features such as formants or pitch. M/EEG-based methods, with their higher temporal resolution, allow us to track how neural responses evolve over time in response to different speech features. Here, we propose a novel EEG-based framework—the Modulation Response Function (MRF)—to examine how cognitive goals shape cortical representations of speech. Building on temporal response function (TRF) modeling, we carefully designed the acoustic predictor to retain fine-grained spectro-temporal modulations critical for speech intelligibility. This tailored representation allows us to reconstruct, from neural responses, a dynamic profile of spectro-temporal encoding in the brain. We then analyze this reconstructed signal in the modulation domain, computing its modulation power spectrum (MPS) to derive the MRF—a cortical analog of the acoustic MPS. This innovative approach enables us to directly investigate whether cognitive tasks (detection vs. comprehension) selectively modulate cortical representations of spectro-temporal features like formants or pitch, which are essential for performing these tasks.

With this approach, we can examine how task-dependent attention shapes cortical representations through the Modulation Response Function (MRF). If attention modulates neural responses uniformly (Gain hypothesis), both tasks should exhibit similar MRF profiles, with any differences reflecting a global spectro-temporal amplification (i.e. gain difference). In contrast, if attention selectively enhances spectro-temporal features based on task demands (Discriminative hypothesis), we expect distinct cortical MRF patterns. For example, cortical responses to formants might be enhanced during comprehension, whereas responses to pitch might be stronger during detection. This would manifest as localized differences in the MRF, with both positive and negative values, indicating task-specific spectro-temporal modulation rather than a uniform shift. Fig 2 illustrates these predicted MRF patterns.

**Fig 2.**
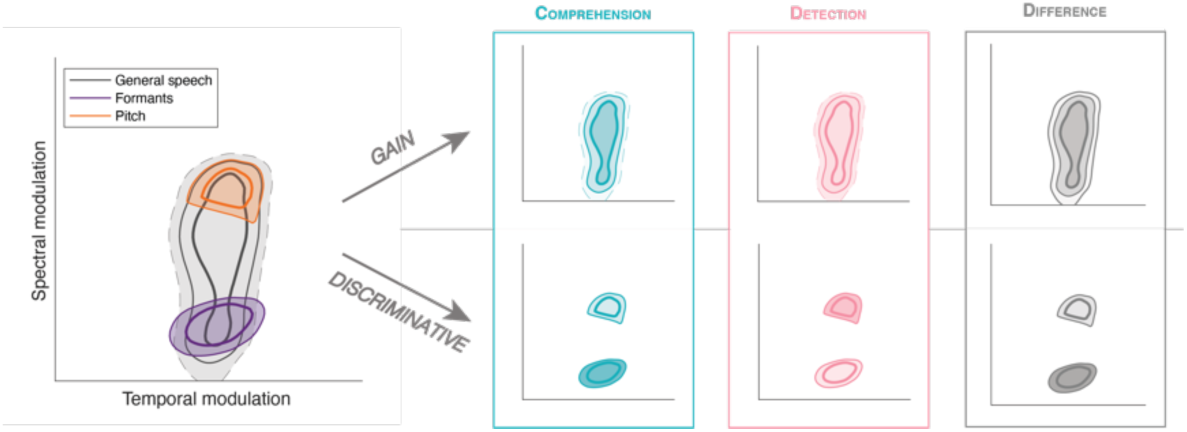
Hypothesis framework. Left panel shows typical spectro-temporal speech features (formants and pitch). According to the Gain hypothesis (upper panels), both cognitive tasks (comprehension (in dark turquoise) and detection (in light pink)) enhance cortical representations, similarly, possibly differing only by global magnitude. Thus, their difference (in gray) would yield uniform positive or negative values across relevant regions. Conversely, the Discriminative hypothesis (bottom panels) predicts distinct cortical filtering of spectro-temporal features depending on the cognitive task. For example, formants may be selectively enhanced during comprehension, while pitch is emphasized during detection, resulting in both positive and negative values in the MRF difference, reflecting specific task-dependent modulations.

## Results

Participants performed both a comprehension and a detection task while EEG was recorded. Both tasks used the same set of auditory stimuli, each combining four overlapping sounds from different categories—including speech, music, and environmental sounds—creating acoustically rich scenes. Speech was present in the stimuli and served as a key target for analysis. In the comprehension task, participants identified a keyword from the spoken sentence; in the detection task, they reported which sound categories were present. Listening difficulty was manipulated by adding background noise that increased cognitive demands without masking critical speech features.

### Behavioral performance

**Fig 3** summarizes participants’ performance across tasks. A linear mixed-effects model (Score ∼ Condition + (1|subject) + (1|trial)) was used to assess main effects. In the comprehension task, no significant general effect of noise condition was observed. However, an analysis by keyword category revealed two significant effects: participants selected the correct keyword more often in the easy condition, whereas semantically related keywords were chosen more frequently in the difficult condition. This shift toward semantic alternatives under challenging conditions suggests that comprehension engages high-level reasoning rather than simple word repetition. Participants did not merely rely on auditory matching but actively processed meaning, indicating a deeper level of cognitive engagement. In contrast, in the detection task, a significant trend emerged, with participants achieving higher accuracy in the easy condition. This effect was primarily driven by the animal and human sound categories, where participants showed better accuracy under the easy condition. No significant differences were observed for the other sound categories (see Table 1). Notably, performance in the detection task approached ceiling for some categories, particularly for narration, which may have limited the ability to detect noise effects in those cases. This ceiling effect likely reflects the overall easier nature of the detection task compared to comprehension.

**Fig 3.**
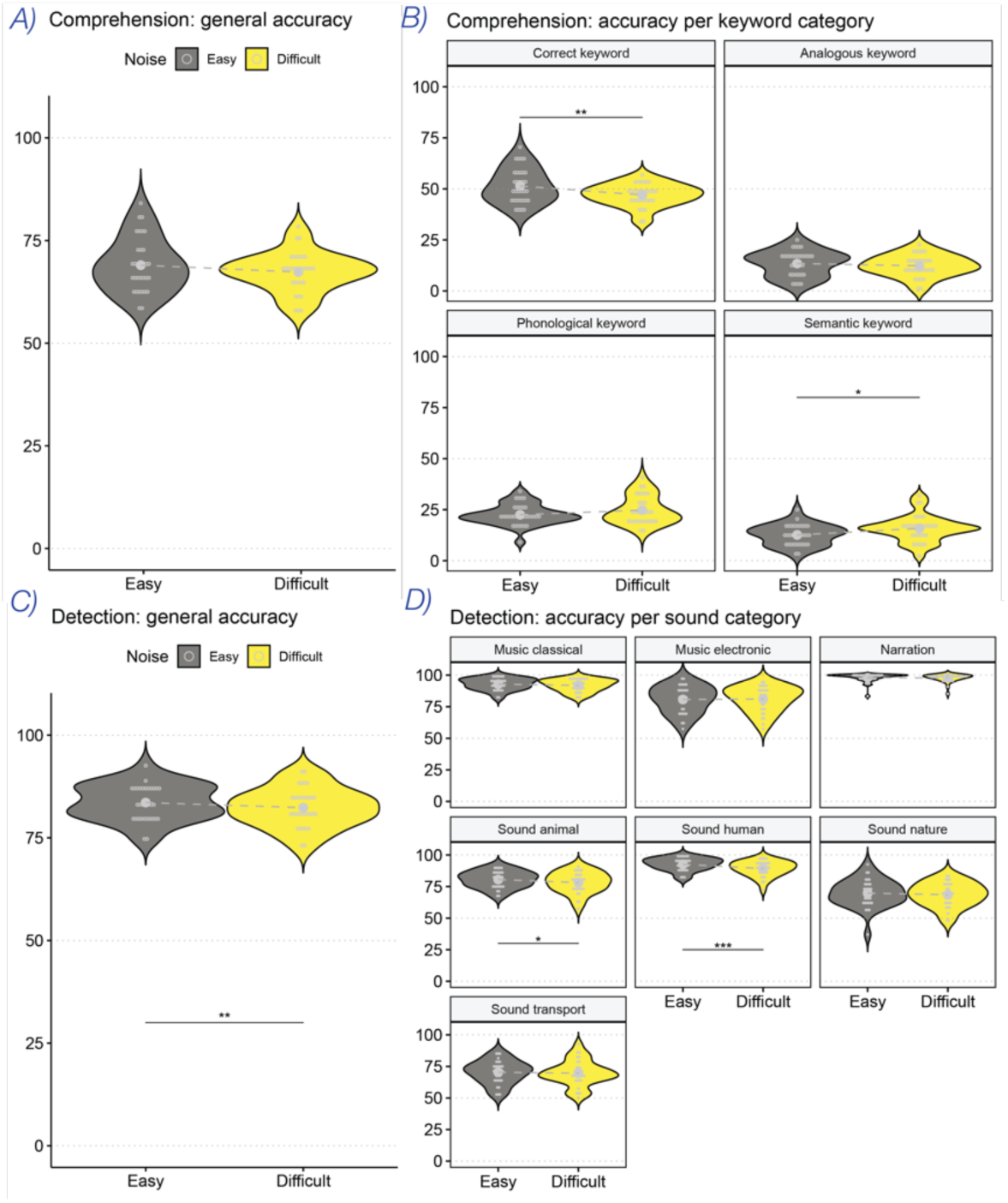
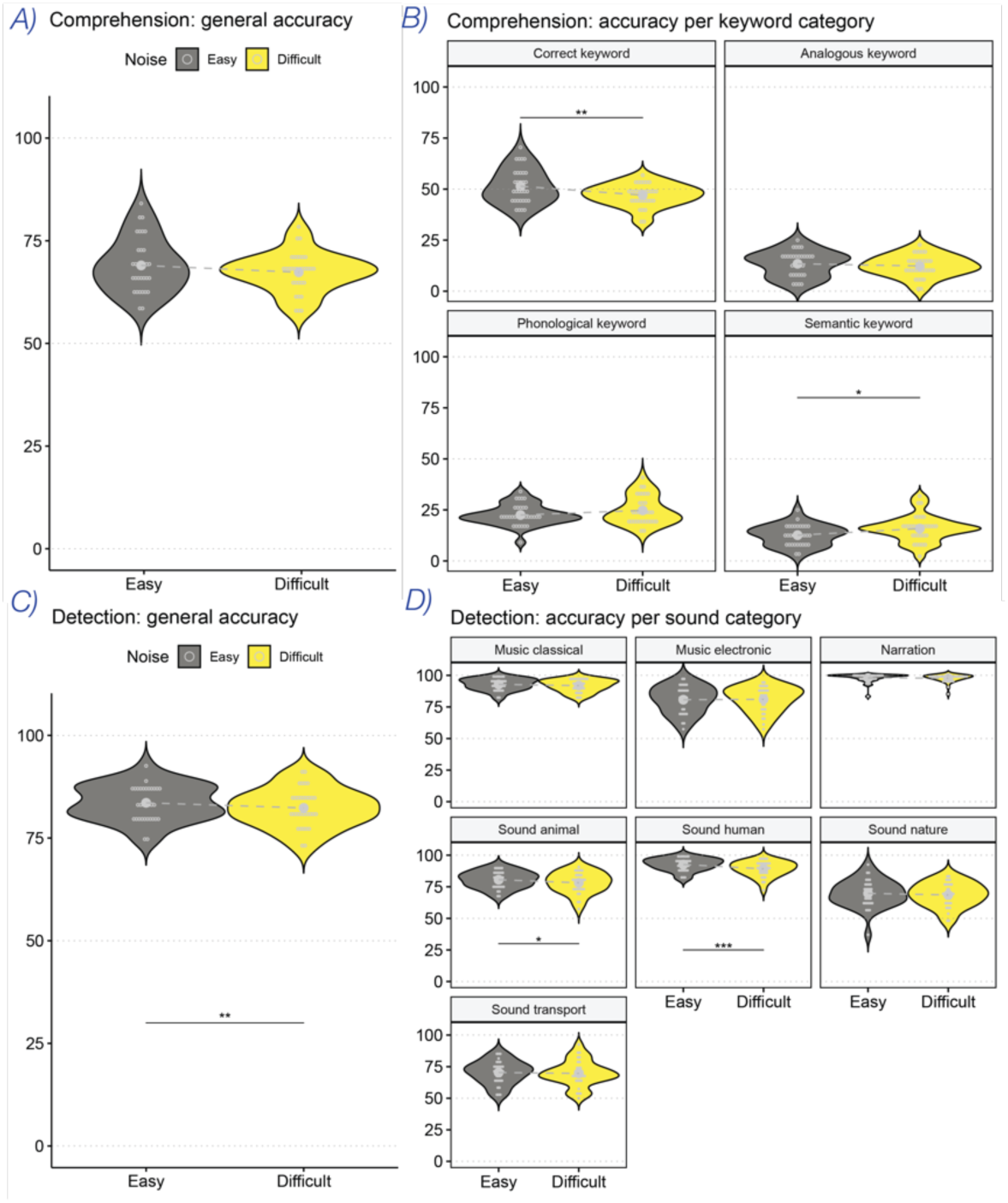
Behavioral accuracy for both tasks. The gray color displays the easy condition while the yellow color is for the difficult condition. Results for comprehension task are shown in panels A (general score) and B (type of keywords). Results for detection task is shown in panels C (general score) and D (type of sound).

**Table 1.** Statistics for behavioral scores.

| Task | Main effect | Statistics |
| --- | --- | --- |
| Comprehension | General | $\chi^2(1) = 2.04, p = 0.15$ |
| | Analogous keyword | $\chi^2(1) = 0.85, p = 0.36$ |
| | Phonological keyword | $\chi^2(1) = 2.32, p = 0.13$ |
| | Semantic keyword | $\chi^2(1) = 6.36, p < 0.05$ |
| | Correct keyword | $\chi^2(1) = 7.08, p < 0.01$ |
| Detection | General | $\chi^2(1) = 6.79, p < 0.01$ |
| | Music: classical | $\chi^2(1) = 1.43, p = 0.23$ |
| | Music: electronic | $\chi^2(1) = 0.03, p = 0.86$ |
| | Narration | $\chi^2(1) = 0.24, p = 0.63$ |
| | Sound: animal | $\chi^2(1) = 4.06, p < 0.05$ |
| | Sound: human | $\chi^2(1) = 14.35, p < 0.001$ |
| | Sound: nature | $\chi^2(1) = 0.81, p = 0.37$ |
| | Sound: transport | $\chi^2(1) = 0.28, p = 0.59$ |

### Neural tracking across tasks

Neural tracking was assessed separately for each task, pooling across noise conditions. A linear mixed-effects model (R ∼ Task + (1|subject) + (1|trial)) revealed a significant effect of task on neural tracking (𝜒^2^(1) = 171.49, 𝑝 < 0.001). Correlations between the reconstructed and original stimulus envelopes (**Fig 4**) were nearly twice as high in the comprehension task (*μ_R comprehension_* = 0.094) compared to the detection task (*μ_R detection_* = 0.047), indicating stronger brain-stimulus synchronization in the comprehension task. Additionally, envelope reconstruction was significantly above chance in both task (*comprehension*: χ^2^(1) = 1860.9, *p* < 0.001; *detection*: χ^2^(1) = 527.43, *p* < 0.001), confirming robust neural tracking in both tasks. Temporal response functions (TRF) analysis (Fig 4) revealed similar components for both tasks, including N100, P200, and N300, while the comprehension task also exhibited a P50. The TRF associated with the detection task appeared flatter, with lower amplitudes across components, suggesting less pronounced neural engagement. Topographical analyses showed activation localized primarily to the left temporal lobe in both tasks, especially around the P200 component. In the comprehension task, additional fronto-temporal activation was observed (linked to the P50), alongside central deactivation corresponding to the N100 (see Supplementary Material for a detailed breakdown of the component and brain area associations). However, these topographical interpretations should be viewed cautiously given the inherent spatial limitations of EEG source localization [45].

**Fig 4.**
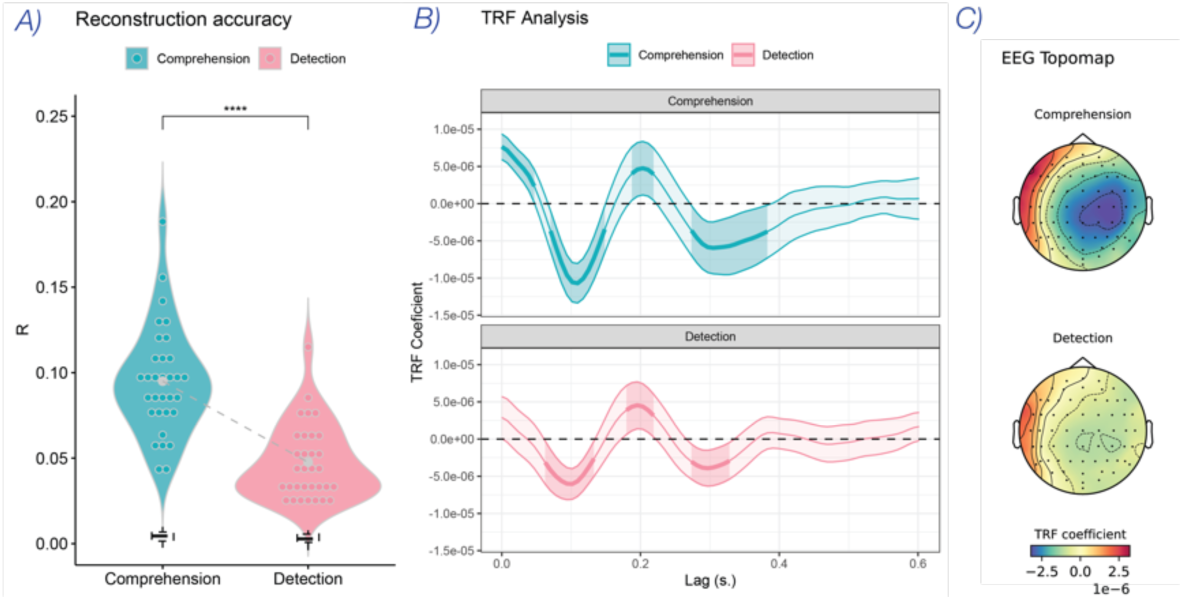
Neural tracking for both tasks. Panel A displays the reconstruction accuracy for the comprehension task in dark turquoise and the detection task in light pink. Each participant’s average is represented by colored round markers, placed within the violin plots, which show the distribution of reconstruction accuracy for each task. The gray central dots represent the grand average per condition, while the dotted line indicates the chance level. The Temporal Response Function (TRF) is represented in panel B for both tasks, using the same color scheme. The shaded ribbons show the confidence intervals, and the bold lines highlight when the TRF significantly differs from zero, marked by the horizontal dotted line. Finally, Panel C shows the EEG topography for both tasks, illustrating the spatial distribution of neural activity.

To test whether selective attention shapes speech representations differently across tasks, we reconstructed the full spectrogram of the speech signals. Unlike envelope tracking, spectrogram reconstruction captures fine-grained acoustic features—such as formants and harmonics—allowing us to examine whether attention enhances these features uniformly (Gain hypothesis) or in a task-specific manner (Discriminative hypothesis; see Fig 2). Separate decoders were trained for each task to reconstruct the speech spectrograms. While overall reconstruction accuracy was lower than for envelope tracking (*μ_R comprehension_* = 0.041; *μ_R detection_* = 0.019), it remained significantly above chance in both tasks(*cpt*: 𝜒^2^(1) = 1910.4., 𝑝 < 0.001; *detect*: 𝜒^2^(1) = 487.3, 𝑝 < 0.001). Critically, these spectrogram reconstructions served as the basis for deriving the cortical Modulation Response Function (MRF), allowing a direct comparison of spectrotemporal tuning across tasks (see Supplementary Material for more information on the spectrograms resconstruction).

MRF profiles for both the comprehension task and the detection task are presented in Fig 5. Despite the relatively low accuracy of spectrogram reconstruction, the modulation profiles still retained a structure similar to that of the original speech signal. Notably, both tasks exhibited a peak around the formants, while a pitch-related peak was more pronounced in the detection task. These patterns suggest that participants focused on the formants during comprehension and on both formants and pitch during detection. A non-parametric permutation test (two-tailed, max-statistic corrected, p<.0001) revealed significant differences between tasks. Specifically, the formant-related peak was significantly stronger in the comprehension task, whereas the pitch-related peak was more pronounced in the detection task. These results align with the Discriminative hypothesis, suggesting that the attentional filter adapts to the specific demands of the cognitive task. In contrast, a Gain hypothesis would have predicted a uniform enhancement of speech-related features favoring one task over the other rather than task-specific modulations. Overall, the cortical MRF magnitude was higher in the comprehension task than in the detection task, consistent with earlier findings of stronger neural tracking during comprehension. To ensure that task-specific differences in spectrotemporal tuning were not confounded by baseline magnitude shifts, we applied a mean-centering procedure across conditions (subjects and trials) prior to statistical testing.

**Fig 5.**
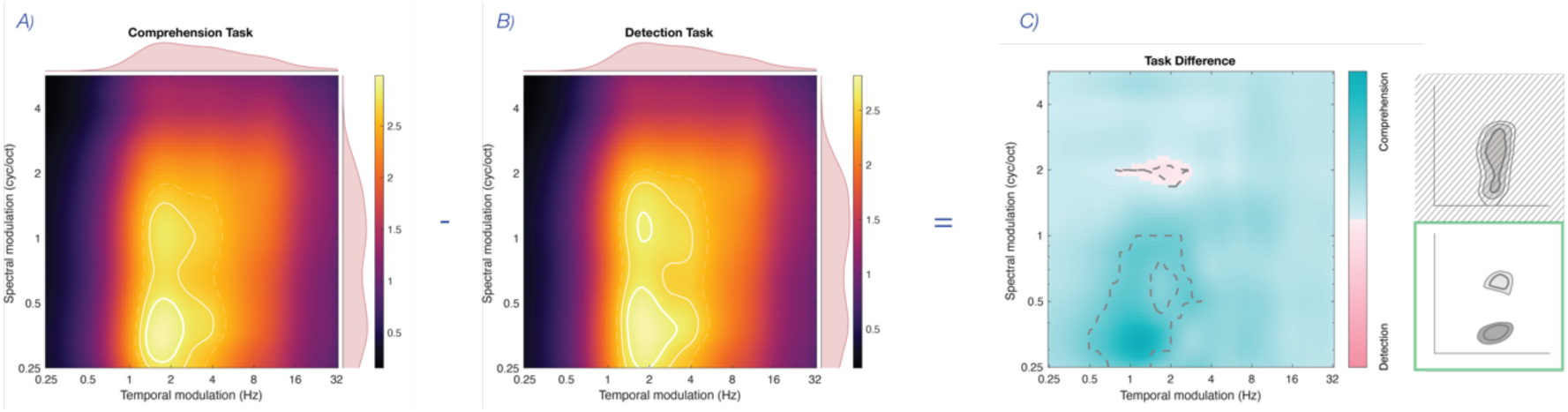
Cortical MRF for both tasks. MRF derived from reconstructed spectrograms in the comprehension task (Panel A). and in the detection task (Panel B). The difference between the two modulation profiles is shown in Panel C. Dark turquoise represents comprehension, and light pink represents detection. Areas within gray dashed lines indicate significant differences computed using a non-parametric permutation test. A schematic representation of the Gain hypothesis is shown in hatching (reproduced from Fig 2); the Discriminative hypothesis is supported, as its predicted pattern more closely resembles the observed significant differences.

Finally, we examined the MRF as a function of participants’ responses in the behavioral comprehension task (see previous section for details). Overall, MRF patterns were similar across response types, with a notable exception: trials where participants selected the phonological keyword exhibited a more prominent peak in the pitch region (Fig 6). Moreover, when directly comparing correct and phonological responses, significant differences emerged in both the pitch and formant regions (𝑝 < .05, 𝑇𝐹𝐶𝐸 − 𝑐𝑜𝑟𝑟𝑒𝑐𝑡𝑒𝑑), with stronger modulation observed for correct keyword selections. Additional comparisons between correct semantic and correct analogous responses also revealed significant differences, although these occurred outside the spectrotemporal regions typically occupied by speech.

**Fig 6.**
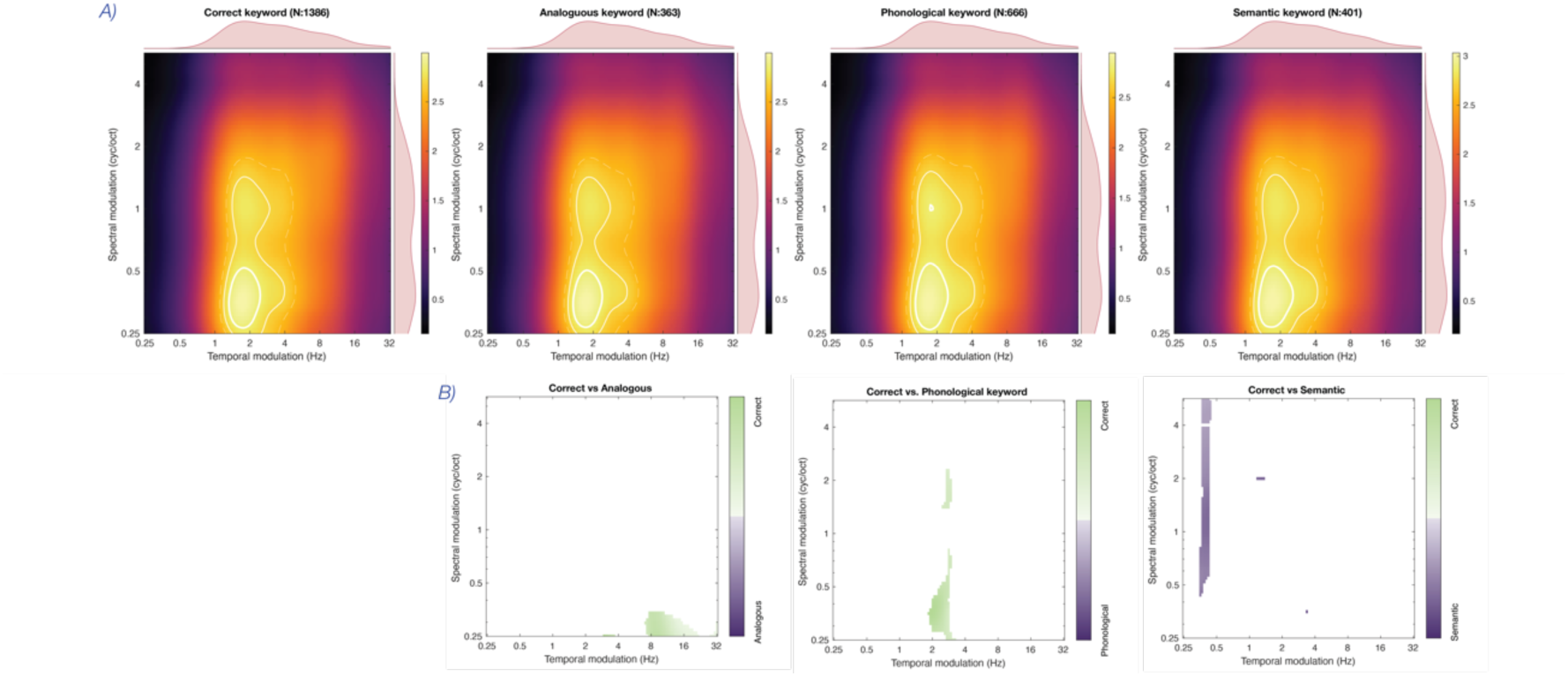
MRF based on participants’ behavioral responses. Panel A shows the cortical MRF associated with participants’ responses. Panel B displays differences in MRFs between trials where the correct keyword was selected (green) and three types of alternative responses (purple): (1) phonologically similar, (2) semantically analogous, and (3) semantically related keywords. Significance was assessed using linear mixed models (LMM) with TFCE correction; only regions showing significant differences are displayed.

### Difficulty condition-specific analysis of neural tracking

In this section, four decoders were trained separately, one for each condition and task, to maximize decoding sensitivity. This approach allows each decoder to optimally capture the relevant neural patterns specific to its associated condition and difficulty. Importantly, control analyses confirmed that the results were similar when a single, shared decoder was applied, ensuring that our findings are not dependent on decoder specificity. A linear mixed-effects model (R ∼ Task * Condition + (1|subject) + (1|trial)) assessed the influence of task and noise difficulty on neural tracking performancxe. No significant effect of noise difficulty was observed, with average correlations remaining similar across conditions within each task (see Table 2 and **Fig 7**), indicating that neural tracking was robust regardless of noise level. Additionally, all the envelope reconstructions were significantly above chance. For the temporal response functions (TRF) analysis (**Fig 7**), TRFs for the comprehension and detection tasks under the easy condition were similar to those described in the previous section, with P50, N100, P200, and N400 components present in both conditions, though the detection task showed lower amplitudes. However, in the difficult condition, TRFs displayed a different pattern: the P200 component disappeared in the comprehension task, replaced by an enlarged N100, while in the detection task, all components were absent except for the N100. For the topographical distribution of neural responses, all conditions showed left temporal lobe activation, primarily associated with the P200. In the comprehension task, additional fronto-temporal activation (linked to the P50) was present under easy conditions, while in difficult conditions, central region deactivation corresponding to the N100 became prominent. (See Supplementary Material for a detailed breakdown of the component and brain area associations).

**Fig 7.**
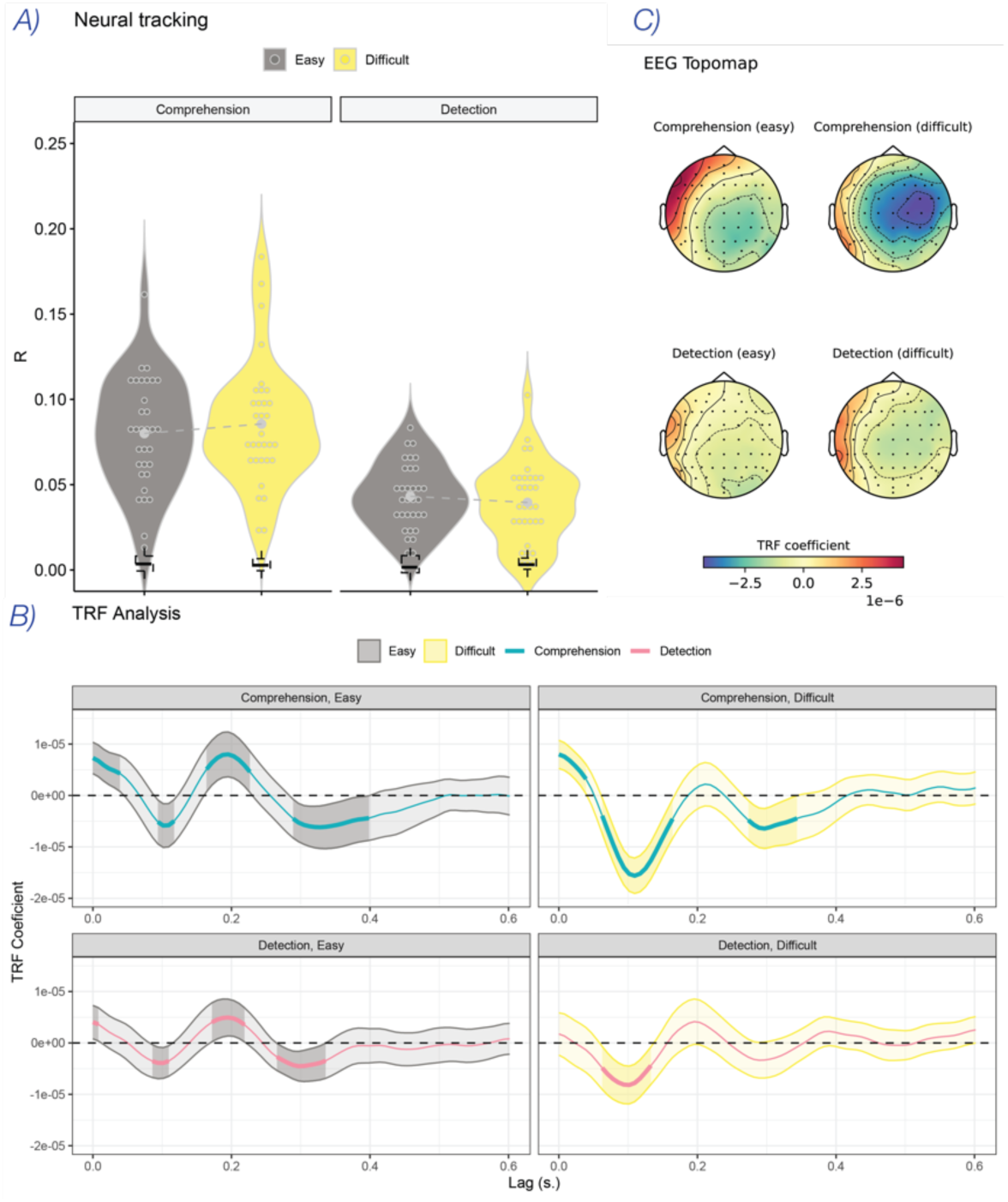
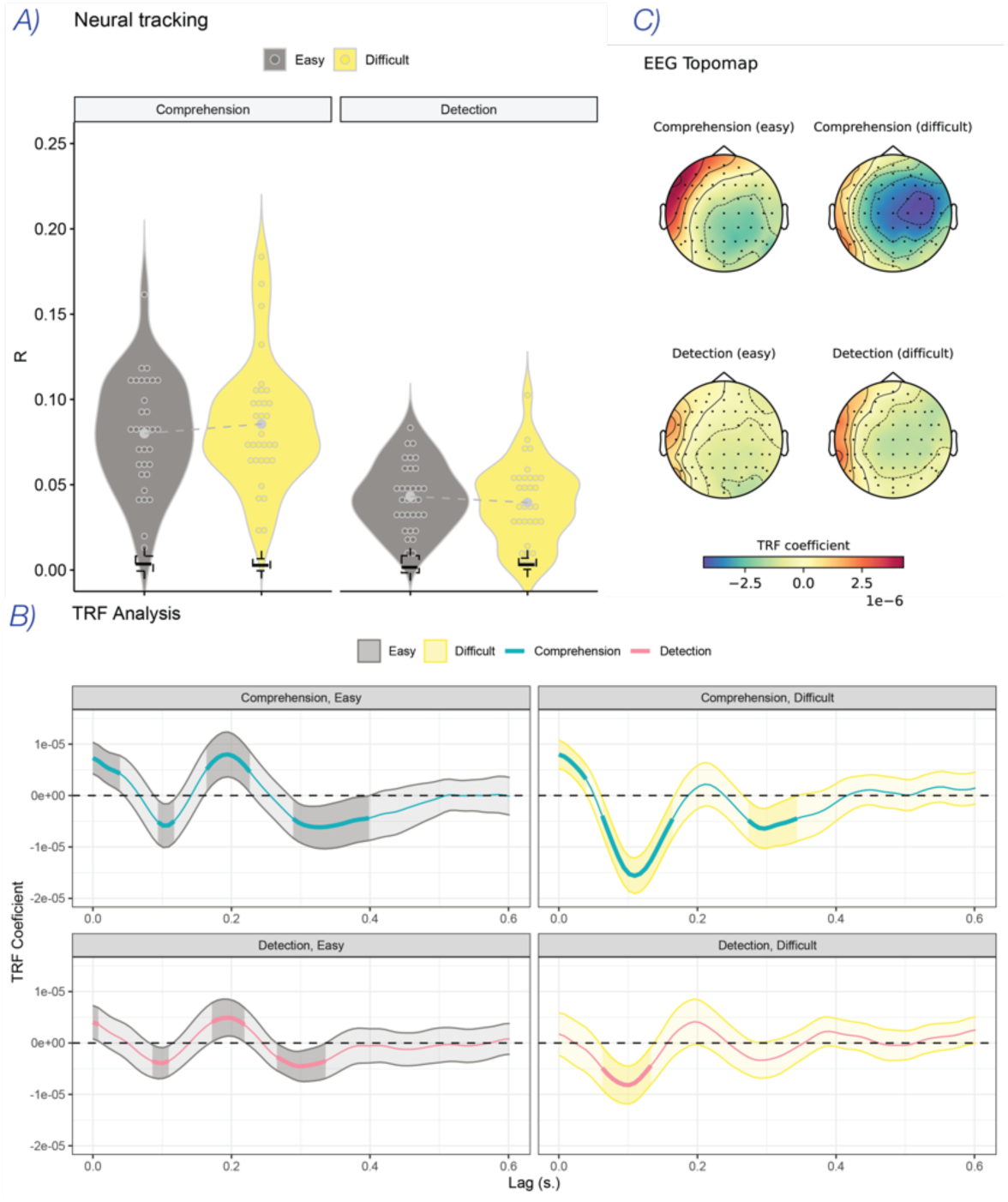
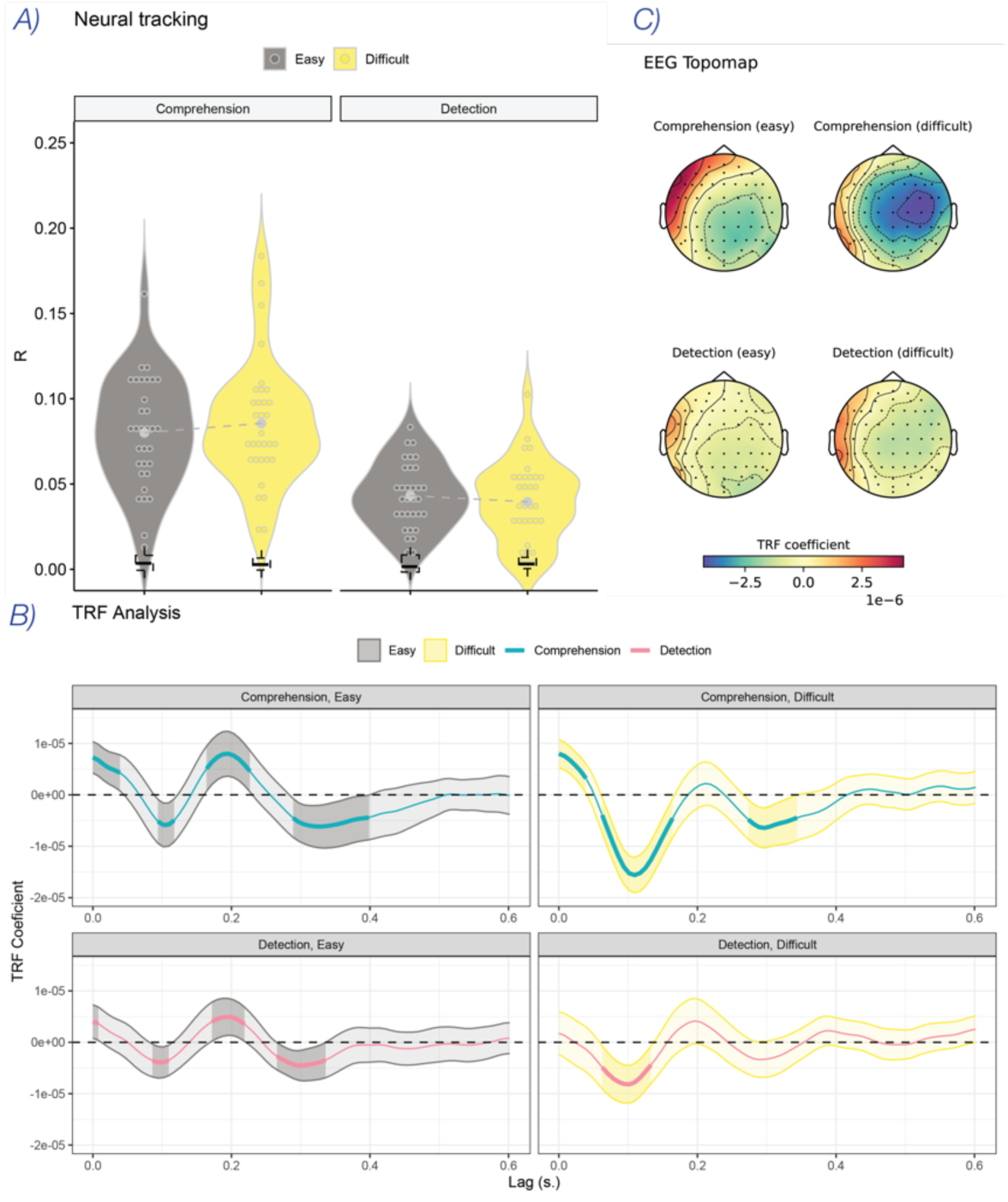
Neural tracking difficulty condition. Panel A) displays the reconstruction accuracy for both tasks for the easy condition in light gray and for the difficult condition in yellow. Each participant’s average is represented by colored round markers, placed within the violin plots, which show the distribution of reconstruction accuracy for each condition. The gray central dots show the average accuracy per condition while the dotted boxplots indicate the chance. The Temporal Response Function (TRF) is represented in panel B for both tasks per difficulty conditions, following the same color scheme. The shaded ribbons represent the confidence intervals, and the bold line highlights the periods where the TRF significantly differs from zero. Finally, panel C show the EEG topography for both tasks under different noise conditions.

**Table 2.** Statistics for the envelope reconstruction for the condition-specific analysis.

| Analysis | Condition | Statistics |
| --- | --- | --- |
| Main effects with LMM | Noise | $\chi^2(1) = 135.44, p = 0.71$ |
| | Task | $\chi^2(1) = 0.14, p < 0.001$ |
| | Task * Noise | $\chi^2(1) = 3.26, p = 0.071$ |
| Condition | Comprehension - Easy | $\mu_R = 0.080; sig: \chi^2(1) = 693.64, p < 0.001$ |
| | Comprehension - Difficult | $\mu_R = 0.086; sig: \chi^2(1) = 749.86, p < 0.001$ |
| | Detection - Easy | $\mu_R = 0.043; sig: \chi^2(1) = 231.55, p < 0.001$ |
| | Detection - Difficult | $\mu_R = 0.040; sig: \chi^2(1) = 172.99, p < 0.001$ |

The spectrograms were reconstructed using four decoders, one for each task and noise difficulty condition. Reconstruction was significant across all conditions (𝑐𝑝𝑡 𝑒𝑎𝑠𝑦: 𝜒^2^(1) = 679.23, 𝑝 < 0.001; 𝑑𝑒𝑡𝑒𝑐𝑡 𝑒𝑎𝑠𝑦: 𝜒^2^(1) = 200.57, 𝑝 < 0.001; 𝑐𝑝𝑡 𝑑𝑖𝑓𝑓𝑖𝑐𝑢𝑙𝑡: 𝜒^2^(1) = 699.08, 𝑝 < 0.001; 𝑑𝑒𝑡𝑒𝑐𝑡 𝑑𝑖𝑓𝑓𝑖𝑐𝑢𝑙𝑡: 𝜒^2^(1) = 158.35, 𝑝 < 0.001), although overall reconstruction accuracy remained low (*μ_R comprehension easy_* = 0.033; *μ_R comprehension easy_* = 0.016; *μ_R comprehension difficult_* = 0.035; *μ_R comprehension difficult_* = 0.015). As shown in Fig 8, under the easy condition, both tasks replicated the MRF patterns observed previously: a clear peak around the formant region for both comprehension and detection, and a more pronounced pitch-related peak for the detection task. In contrast, under the difficult condition, the modulation profiles diverged: difficult comprehension elicited a new peak in the pitch region, while difficult detection exhibited a broader enhancement encompassing both formant and pitch regions. Across-task comparisons revealed that under easy listening conditions, participants primarily enhanced formant cues during comprehension and pitch cues during detection (p < .01, permutation test, baseline corrections were applied as described in the Methods). In the difficult condition, however, the formant region showed significant differences between the two tasks, with increased engagement observed during detection. Taken together, these patterns suggest that under easy listening conditions, attention selectively enhanced formant cues for comprehension and pitch cues for detection. Under more challenging conditions, attentional focus appeared to shift: comprehension began to incorporate pitch-related information, while detection recruited a more distributed spectro-temporal representation spanning both formants and pitch. Within-task comparisons further support this interpretation. In the comprehension task, formant-related enhancement was broader under easy conditions than under difficult ones, suggesting that increased listening difficulty may reduce reliance on mid-frequency spectral cues. In the detection task, a similar—though more modest—difference in formant-related activity was observed, again pointing to a redistribution of attentional resources under more challenging conditions.

**Fig 8.**
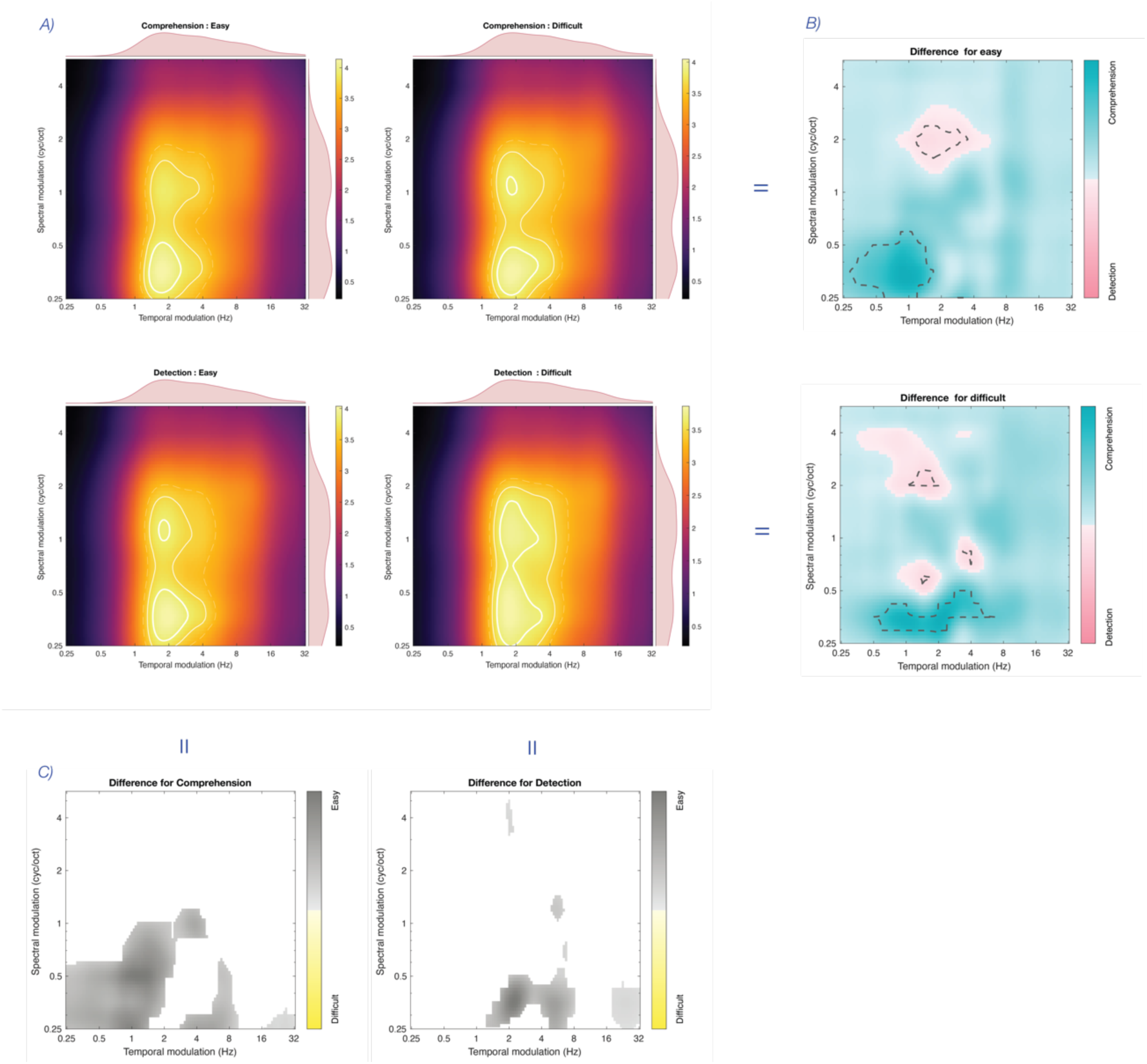
MRF by difficulty noise. Panel A shows the MRF for the four experimental conditions: easy comprehension, difficult comprehension, easy detection, and difficult detection. Panel B illustrates the difference between the two tasks for each difficulty condition, with dark turquoise representing comprehension and light pink representing detection. Areas within gray dashed lines indicate significant differences computed using a non-parametric permutation test. Panels C displays the difference between the difficulty conditions for each task, with gray representing easy condition and yellow representing the difficult condition. Significance was assessed using linear mixed models (LMM) with TFCE correction; only regions showing significant differences are displayed.

## Discussion

### Cognitive goals flexibly reshape sensory encoding

Selective attention is often conceptualized as a mechanism that enhances the neural encoding of relevant stimuli. Yet a critical question remains: does attention amplify all features of a selected sensory input equally, or does it adaptively reshape sensory encoding based on cognitive goals? The current study addressed this question by examining how cortical representations of speech are modulated when participants engage in either a comprehension task or a detection task, using identical acoustic stimuli.

Across both tasks, behavioral performance was high. In the comprehension task, participants consistently selected the correct keyword over distractor keywords. Performance slightly declined in the more difficult noise condition, and notably, semantic keywords were chosen more frequently in this condition. This suggests that when comprehension became more effortful, participants may have relied more heavily on top-down semantic cues. In the detection task, overall accuracy remained stable across noise difficulty for most sound categories. However, under more difficult noise, accuracy decreased specifically for human and animal sounds. Crucially, detection performance for the narration stream—the speech content used for EEG analysis—remained near ceiling across all conditions, confirming that participants remained engaged and able to process speech even in more challenging acoustic environments.

At the neural level, our findings reveal that selective attention does not act as a uniform amplifier across speech features, but instead dynamically modulates spectro-temporal representations based on task demands. Envelope reconstruction accuracy was stronger in the comprehension task, indicating greater cortical tracking of speech when semantic processing was required.. Cortical Modulation Response Function (MRF) further revealed that different spectro-temporal features were selectively enhanced depending on cognitive goals: during comprehension, formant-related modulations —critical for phonemic decoding—were amplified, whereas during detection, pitch-related modulations —essential for speaker and speech presence detection—were more prominent. These patterns were further modulated by noise difficulty. Under difficult noise, spectro-temporal encoding patterns shifted, further emphasizing the dynamic nature of auditory attention.

These results allowed us to adjudicate between two competing models of attentional modulation: the Gain hypothesis, which posits a uniform amplification of all features of the attended input, and the Discriminative hypothesis, which proposes a goal-sensitive, feature-specific enhancement. Our findings more clearly support the Discriminative hypothesis. The task- specific spectral tuning—favoring formants for comprehension and pitch for detection—demonstrates that attention adaptively prioritizes the features most relevant for the current cognitive objective, rather than globally amplifying the entire speech signal.

While overall MRF magnitude was higher during comprehension, we do not interpret this as strong evidence for a general gain mechanism. Instead, we suggest it more likely reflects heightened listening effort and cognitive engagement. Listening effort refers to the increased cognitive resources required to process speech under challenging conditions [46].

Supporting this view, neural tracking has been shown to reflect not only speech intelligibility but also the listener’s cognitive engagement. For instance, neural tracking increases with listening difficulty—even when behavioral performance remains stable—suggesting that it captures sustained attentional effort under adverse conditions [47–49]. In line with this interpretation, we found that neural tracking significantly predicted overall MRF magnitude, suggesting a link between sustained attentional encoding and spectro-temporal enhancement at the cortical level. In our study, although neural tracking was consistently stronger for comprehension than for detection, we did not find evidence of similar cortical magnitude increases between easy and difficult noise conditions within each task, which may indicate that the neural effects of noise were too subtle to be captured, even though behavioral measures showed sensitivity to noise difficulty.

Importantly, our findings show that attention flexibly reconfigures sensory encoding within a fixed sensory domain— speech—rather than merely selecting between different sensory streams. Although both tasks used identical speech stimuli, cortical responses were distinctly shaped by whether the listener aimed to detect the presence of speech or comprehend its meaning. This task-dependent reconfiguration of sensory encoding underscores the flexibility of attention as an adaptive mechanism, aligning neural representations with cogntive goals. In sum, selective auditory attention emerges not as a passive gain control mechanism, but as an active, adaptive process that flexibly reshapes cortical sensory encoding to align with shifting cognitive goals.

### Exploratory insights into neural representations of speech

To further examine how these attentional modulations relate to behavior, we examined trials from the comprehension task based on participants’ keyword choices. Correct responses were associated with stronger cortical modulations in both pitch and formant regions, suggesting more accurate encoding of its spectro-temporal features. In contrast, trials where participants selected the phonological keyword showed enhanced pitch-related modulations. This may reflect a less precise encoding of the speech signal, leading listeners to rely on broad acoustic similarity (e.g., pitch contour) rather than detailed phoneme-level information. Semantic keyword, also absent from the stimulus, were more frequently selected under difficult noise, yet their associated MRF profiles did not significantly differ from correct trials. This may indicate that semantic interpretations were constructed independently of strong acoustic cues, or that such high-level processes, while potentially present, remain undetected due to the MRF’s limited sensitivity to abstract linguistic encoding. This dissociation aligns with previous findings showing that semantic dissimilarity is tracked by later and more distributed neural responses [50]. Together, these results suggest that while MRF reflects auditory encoding, behavioral responses also capture interpretive processes that unfold beyond early acoustic representations.

While MRF captures the spectral structure of encoded speech, we also examined the temporal dynamics of neural responses using TRF-based analyses to gain insight into the temporal dynamics of processing. Although TRF analyses offer valuable insight into the temporal dynamics of speech processing, their interpretation requires caution due to methodological challenges and inter-individual variability [51,52]. In the present data, we observed P50–N1–P2 components [48] consistent with early sensory encoding and attentional modulation of speech features. The P2 component, typically linked to perceptual categorization, was only observed in easier conditions, and disappeared under high difficulty—likely reflecting a disruption of mid-level auditory processing. In addition, a central negative component peaking around 300 ms was observed exclusively in the comprehension task, across both easy and difficult conditions. Although this deflection does not match the typical characteristics of the semantic N400 observed in studies using linguistic predictors [50], its latency is consistent with the N300, which has been linked to phonological expectancy and categorical priming [53]. Its selective emergence during comprehension, but not detection, suggests that comprehension engages additional, task-specific linguistic processes beyond low-level auditory monitoring. These findings further support the idea that cognitive goals flexibly shape the cortical encoding of speech.

### Methodological insights and future perspectives

Methodologically, this study introduced several key innovations. Foremost among them is the development of the cortical Modulation Response Function (MRF), derived from spectrograms reconstructed from EEG data. To our knowledge, this is the first instance in which EEG-based spectrogram reconstructions—despite their relatively low absolute accuracy— have been leveraged to extract meaningful spectro-temporal representations of cortical processing. Remarkably, these reconstructions, although coarse, retained task-relevant structure and revealed interpretable patterns of spectro-temporal encoding. This novel use of the MRF framework enables a more direct and fine-grained analysis of how specific acoustic features are neurally represented.

A second key innovation is our approach to designing the noise. Unlike classical energetic masking, which broadly disrupts the signal by overlapping spectrally or temporally [54], or informational masking, which introduces linguistic or perceptual competition [55], our approach strategically avoided overlapping with speech-critical modulation regions. By shaping the noise in modulation space, we were able to systematically vary task difficulty while preserving the intelligibility of the speech signal. This method offers a more precise way to investigate how noise and cognitive demands interact in shaping neural representations. Additionally, by using identical acoustic stimuli across tasks, we ensured that the results we observed were driven by the nature of the cognitive task itself—detection versus comprehension—rather than differences in the stimuli.

Nevertheless, one limitation concerns the task structure: both employed closed-set response formats, but only the detection task provided participants with prior knowledge of the possible response categories. This asymmetry may have influenced the attentional strategies available to them. In detection, listeners could pre-allocate attention to known targets, whereas in comprehension, they needed to attend broadly and flexibly to the entire narrative. Prior knowledge has been shown to facilitate neural speech encoding and enhance tracking of relevant speech streams, particularly by modulating cortical responses through expectation-driven mechanisms [56,57]. This potential difference may have contributed to the stronger tracking observed in comprehension and should be carefully considered in future designs.

Another potential methodological limitation relates to the spectral characteristics of the reconstructed spectrograms. Specifically, we observed a shift of temporal modulation frequencies toward lower values (∼1–2 Hz) in the MRF. This may partly result from the regularization applied during TRF estimation, which smooths the TRF, attenuating rapid temporal fluctuations [52]. Alternatively, this shift might reflect intrinsic properties of cortical auditory processing, which preferentially integrates acoustic signals at slower temporal scales, corresponding notably to lexical and phrasal structures occurring around 1–2 Hz, as opposed to faster syllabic modulations typically observed around 4–7 Hz [58]. In our study, this bias towards slower temporal modulations may reflect the demands of the cognitive tasks, which required participants to extract meaning or identify speech within noisy scenes. Such tasks and noise difficulty may naturally emphasize processing at the lexical level, rather than at the syllabic rate, particularly when higher-level linguistic goals are prioritized. These methodological and neurophysiological factors are not mutually exclusive, and future research should account for both when interpreting modulation-based neural representations.

While envelope and spectrogram reconstruction remain useful tools to assess neural tracking, they are constrained by the limitations of time-frequency space. By contrast, MRF captures neural sensitivity across spectro-temporal modulation dimensions—features that are critical for speech intelligibility and ecological listening. Crucially, this approach is not limited to speech or specific task paradigms. It can be extended to a broad range of auditory environments and may be combined with model-based decoding techniques—such as spectro-temporal receptive fields or neural networks—to quantify how attention dynamically reshapes cortical tuning across complex sensory scenes.

## Materials and methods

### Stimuli design

To test our hypothesis, we designed auditory stimuli suitable for both comprehension and detection tasks. These stimuli were composed of four sound samples (or stems), each drawn from one of seven predefined sound categories and distributed throughout the duration of the stimulus.

#### Sound categories

We largely grounded our categorization of stimuli in seven distinct sound types based on established classifications of sounds [59,60], namely: narrative stories, two types of music, and four categories of environmental sounds. For the narrative stories, we extracted short excerpts from three audiobooks, all narrated by the same male voice, Scott Brick, characterized by a deep pitch and controlled prosody [61–63]. These excerpts were selected because they functioned as standalone content, maintaining coherence even when removed from their broader context. For the musical stimuli, we selected two types of music: slow-tempo classical pieces (70–85 BPM), and fast-tempo electronic music (115–140 BPM). The tracks were sourced from SoundCloud and the Free Music Archive [64,65], and their tempos were analyzed using madmom and librosa [66,67]. In addition to the narration and music, we curated a diverse set of environmental and non-verbal human sounds from various online sources [68–70]. These included:

- Animal sounds (e.g., donkey, pig, hens, dog, birds)
- Non-speech human sounds (e.g., laughter, moans, screams, breathing, crying)
- Nature sounds (e.g., wind, sea, rain, thunder, waterfalls)
- Transport sounds (e.g., car, train, bus, jet, helicopter)

All sound samples were carefully selected for high audio quality, then normalized and trimmed to a duration between 7 and 11 seconds. Notably, all sound categories occupied distinct regions in the modulation profile, as illustrated in Fig 9.

**Fig 9.**
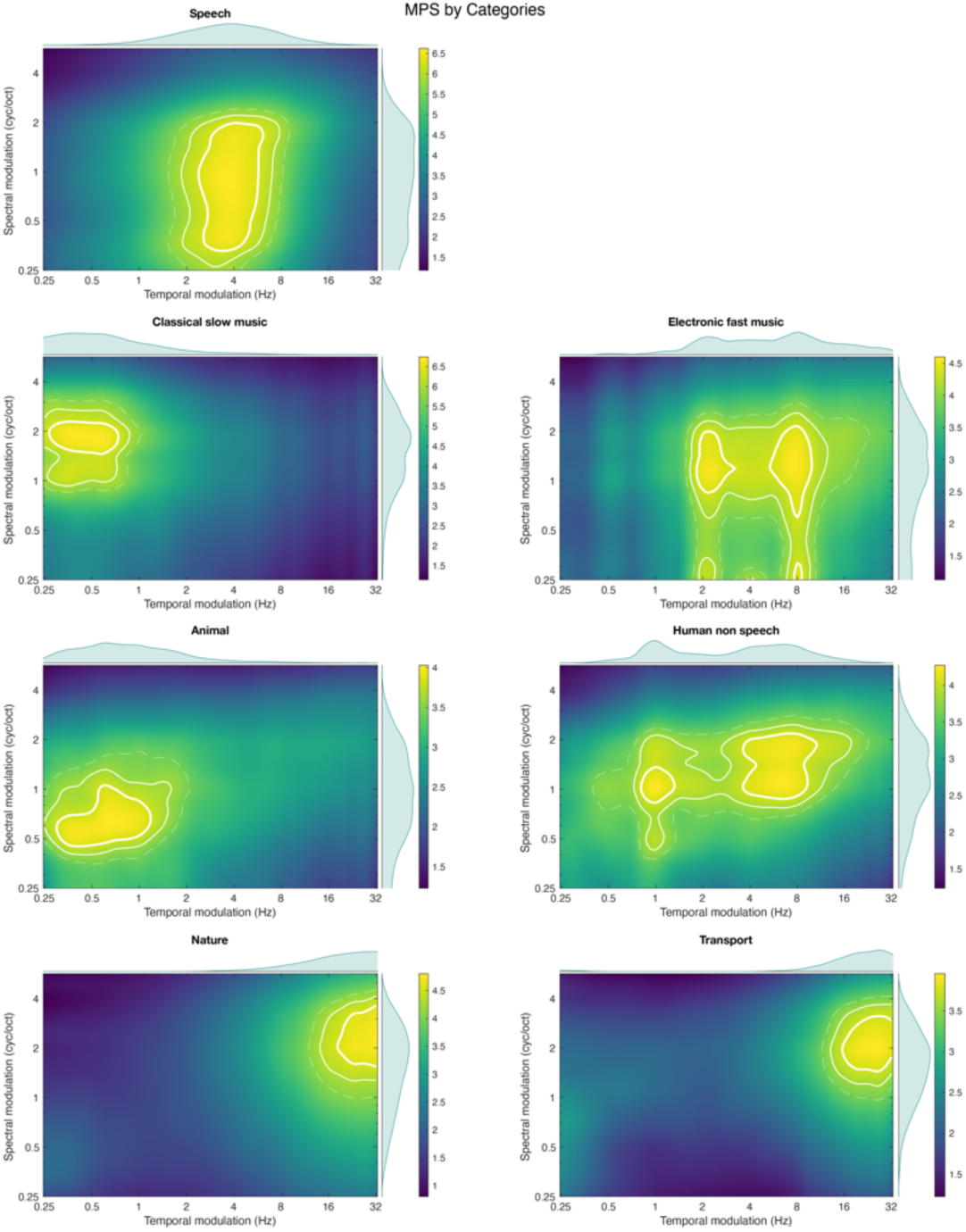
Acoustic MPS by categories. Each sound category is represented by its modulation profile, with temporal modulation (in hertz) on the x-axis and spectral modulation (in cycles per octave) on the y-axis. The bold white line marks the 95^th^ percentile, the solid white line marks the 90^th^ percentile, and the dashed white line marks the 85^th^ percentile. The average distribution along the spectral modulation axis is displayed to the right of each profile, while the average distribution along the temporal modulation axis is shown above each profile.

#### Stimuli composition

Each stimulus consisted of four different sounds selected from distinct categories (classical music, electronic music, animal sounds, nature sounds, human sounds, transport sounds, and narration). The total duration of each stimulus was 15 seconds, sampled at 8 kHz with 16-bit resolution (uncompressed, mono). The four sounds were arranged in different predefined sequences so that the order varied across trials, ensuring that the same categories did not always appear first or last. To introduce variability and avoid systematic overlaps, each music category (classical or electronic) appeared in 40.9% of the stimuli, and each environmental sound category appeared in 54.5% of the stimuli. Additionally, 31.8% of the stimuli contained no music at all, and 13.6% contained no environmental sounds. When narration (speech) was included, its position was chosen from predefined sequences to ensure a balanced sound distribution and limit category overlap within the 15-second window. Since every narration contained a target keyword, its timing guided sequence selection. The optimal configuration minimized keyword overlap with other sounds, ensuring it was not masked by more than two and occurred neither too early nor too late. To create smooth transitions, a 750ms fade-in was applied at the beginning of each sound and a fade-out of the same duration at the end. This prevented sudden onsets and offsets, making the transitions more natural and reducing abrupt perceptual changes that could draw unnecessary attention to specific sounds. Finally, a specifically designed noise was added to the entire stimulus at a signal-to-noise ratio (SNR) of 2 dB. The resulting stimulus set thus provided a controlled but variable acoustic environment, with balanced representation of different sound categories across trials.

#### Noise design

To manipulate task difficulty while preserving speech intelligibility, we designed specific noises using the cortical representation functions from NSL Toolbox [71]. This process involved generating a mask template that defined the ideal spectral and temporal distribution of the noise within the modulation profile (rate and scale domains). The mask was designed to be anti-speech, meaning that it was selectively applied only to the regions of the MPS not associated with key speech modulations. This ensured that the added noise avoided direct overlap with key speech modulations, thereby maintaining speech intelligibility while still introducing task-related interference (see Supplementary Material for details). To systematically vary task difficulty, we adjusted the mask size, with larger masks covering more of the modulation profile and increasing interference, making the task more challenging. Smaller masks preserved more of the speech signal, resulting in clearer speech and an easier task. Once the mask was defined, it was applied to the modulation representation of white noise, shaping its spectro-temporal structure. The noise was then reconstructed. To determine the optimal mask parameters, we conducted preliminary studies (see Supplementary Material for details). In this experiment, we used two types of noise: an "easy" noise, which remained separated from speech regions in the modulation profile, and a "difficult" noise, which overlapped with speech regions, thereby increasing interference (as illustrated in Fig 10).

**Fig 10.**
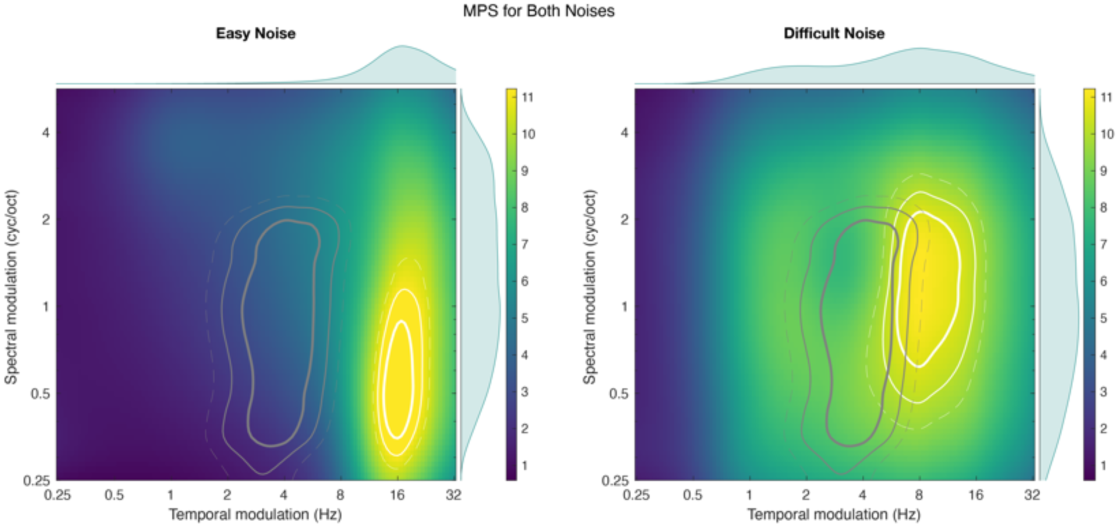
Acoustic MPS for both noises. For each noise sound, the modulation profile is represented with temporal modulation (in hertz) on the x-axis and spectral modulation (in cycles per octave) on the y-axis. The bold solid line indicates the 95^th^ percentile, the solid line indicates the 90^th^ percentile, and the dashed line indicates the 85^th^ percentile. White lines correspond to the modulation profile of the white noise, while dark gray lines represent the modulation profile of the speech signal. Notably, the speech profile shows greater overlap with the difficult noise than with the easy noise.

### Experimental design

Participants completed two tasks — comprehension and detection — using a shared core set of stimuli, allowing for a direct comparison of the two cognitive processes. The primary difference between the tasks lay in the cognitive demands they imposed. To control for potential order effects, the sequence of tasks was counterbalanced across participants. Each stimulus was presented in either an easy or a difficult noise condition, resulting in a 2 (task: comprehension vs. detection) × 2 (noise difficulty: easy vs. difficult) design with an equal distribution of conditions.

#### Detection Task

In the detection task, participants identified the different sound categories present in each stimulus among seven predefined categories. At the end of each stimulus, the names of all seven categories were displayed simultaneously, and participants could select as many categories as they believed had been present. This closed-set, multiple-response format ensured that participants had full knowledge of the possible options but not of the number of correct answers on each trial.

To prevent participants from relying solely on the presence of narration to perform the task, a subset of stimuli without narration was included. As a result, the proportion of stimuli containing narration was reduced to 81.5%. Despite this adjustment, the relative distribution of the other sound categories was preserved, with each music category present in 44.4% of the stimuli and each environmental sound category present in 57.4%. Additionally, 27.8% of the trials contained no music, and 8.3% lacked environmental sounds. This design maintained category balance while introducing variability in the acoustic composition of the stimuli.

### Comprehension Task

In the comprehension task, participants focused solely on the narration and, at the end of each stimulus, selected the correct keyword uttered by the narrator from a set of four options. This closed-set, post-stimulus word recognition format is similar to that used in a previous study, where participants engaged in a forced-choice comprehension task [72]. For the comprehension task, each original keyword, extracted from the speech narration, was systematically paired with three related words based on close phonological and semantic relationships. A semantic distractor is selected for its strong semantic similarity to the original keyword, whereas a phonological distractor is identified based on its phonological resemblance to the original keyword. An analogical distractor is derived by ensuring it is semantically related to the phonological keyword and phonologically similar to the semantic keyword. For example, in the sentence, *"Scientists had always classified dinosaurs as reptiles, cold blooded creatures drawing the heat they needed for life from the environment."* the original keyword *"cold"* is paired with:

- *"hold"* as the phonological keyword, sharing similar sounds with "cold".
- *"cool"* as the semantic keyword, being conceptually related to the idea of temperature.
- *"keep"* as the analogical keyword, bridging the semantic relationship with "hold" (both imply retention) and the phonological similarity with "cool" (short vowel and consonant pattern).

See Supplementary Material for more details on the selection of keywords. This structured selection process establishes an interconnected framework of phonological and semantic associations, enabling a nuanced investigation of their interplay in language processing and cognitive mechanisms.

#### Procedure

The experiment consisted of eight stimulus blocks for the comprehension task and eleven blocks for the detection task. Before each task, participants completed a training block to familiarize themselves with the procedure. Between blocks, they had self-paced breaks, allowing them to resume the experiment whenever they were ready. The session took place in a sound-attenuated booth, where participants listened to stimuli through Etymotic ER-2 earphones. At the beginning of the session, they were instructed to adjust the volume to a comfortable level. Stimuli were presented using OpenSesame [73]. To minimize artifacts in the electroencephalographic (EEG) recording, participants were instructed to avoid eye movements throughout the experiment. Data collection lasted between 70 and 115 minutes, and the entire procedure was completed in a single session. All procedures were approved by the Institutional Review Board (IRB).

### Participants

Thirty-two participants, aged between 18 and 30 (μ = 20.4 years, σ = 2.2), took part in the experiment. All participants were undergraduate or graduate students and reported no hearing nor cognitive impairments. They provided informed consent before participating in the study and received research credit for their participation.

### EEG acquisition and signal processing

Electroencephalographic (EEG) data were recorded using a BioSemi system with 64 channels at a sampling rate of 2048 Hz. Signal processing was performed using the MNE-Python framework [74]. The preprocessing pipeline included downsampling to 128 Hz, band-pass filtering between 2 and 30 Hz, and detection and interpolation of noisy channels using PyPREP [75].

The stimulus envelope was extracted using a gammatone filterbank spanning 125 Hz to 8000 Hz. The envelope was computed by taking the absolute value of the filtered signal, raising it to the power of 0.6, and averaging across filter outputs. The resulting envelope was then resampled to 128 Hz for further analysis.

The spectrogram was generated using a cochlear gammatone filterbank, decomposing the signal into logarithmically spaced frequency bands between 125 Hz and 8000 Hz, with 32 filters per octave. Each band underwent half-wave rectification and smoothing, reducing noise while preserving critical signal features. The spectrogram was then logarithmically compressed to enhance its dynamic range and resampled to 128 Hz to match the EEG data sampling rate.

### Data analyses

#### Backward modeling and stimulus reconstruction

To investigate the relationship between neural activity and stimulus features, we used regularized linear regression to reconstruct both the speech envelope and the spectrogram from EEG recordings. The decoders were computed using the MNE- Python library following a backward modeling approach, in which neural responses are mapped back to stimulus features. These decoders, conceptually equivalent to backward TRFs, consist of sets of weights estimated across 𝑁 electrodes and different time lags 𝜏. Here, we considered time lags from -600 to 0ms. For a given trial, the stimulus reconstruction 𝑆^_^(𝑡) was computed as:

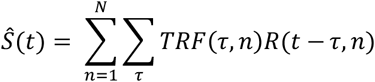

Where 𝑅 represents the shifted neural responses from each electrode 𝑛 at time 𝑡 = 1. . 𝑇. The backward TRF was estimated using ridge regression, formulated as:

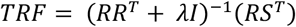

where 𝜆 is the regularization parameter, chosen to optimize the reconstruction, and 𝐼 is the identity matrix. The optimal ridge parameter was set 𝜆 = 10^=^ according to Crosse et al., 2016.

TRFs were estimated on a trial-by-trial basis for each subject and task. Stimulus reconstruction was performed using a leave-one-out cross-validation approach: each trial was reconstructed using a TRF trained on the 87 remaining trials, ensuring that the test trial was never included in its own training set. Since each task contained 88 speech trials, this process was repeated for every trial. Additionally, the same procedure was applied at the level of task difficulty, where trials were grouped accordingly, resulting in 44 trials per condition and a total of four distinct TRFs. Reconstruction accuracy was assessed using the Pearson’s correlation coefficient between the reconstructed and original signals. Finally, to confirm that the observed effects were not due to random chance, each reconstructed signal was also compared to other speech signals as a control measure. This control measure ensured that the observed reconstruction accuracy reflected genuine speech tracking rather than incidental correlations with any speech signal.

#### Gabor modulation profile

To analyze both the acoustic modulation power spectrum (MPS) and the cortical Modulation Response Function (MRF), we employed Gabor filters to extract the spectrotemporal modulation profile of the auditory signal. The modulation profile can be computed using various techniques, depending on the specific objectives of the analysis [76]. The signals (original spectrogram or reconstructed spectrogram from TRF) were processed through a series of transformations to derive the modulation profile. The spectrogram 𝑦(𝑡, 𝑓) was analyzed using two-dimensional Gabor filters to capture the spectrotemporal characteristics of the sound [77]. Each filter was tuned to a particular temporal modulation (or rate) 𝜔 in Hz and a spectral modulation (or scale) Ω in cycles/octave. The Gabor filter had specific Gaussian bandwidths to capture rapid changes and was defined as follows:

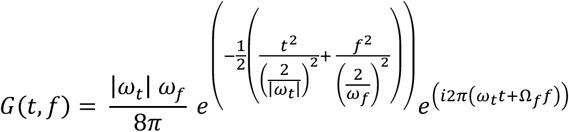

Finally, the auditory spectrogram was convolved in both time and frequency with the Gabor filters, yielding a projection into modulation space. The resulting four-dimensional output R was collapsed along the time and frequency axes to create the rate-scale modulation profile. Additionally, positive and negative temporal rates were also collapsed to average the modulation profile across both directions.

#### Response coding

In the comprehension task, participants identified the keyword spoken by the narrator from a set of four options. Accuracy was scored using a graded system: 1 point was awarded for selecting the correct keyword, 0.5 points for selecting a phonologically or semantically related keyword, and 0 points for selecting the analogous keyword. This scoring approach allowed us to capture partial recognition performance and provided a more nuanced measure of speech comprehension than binary accuracy.

#### Statistical analyses

We used (generalized) linear mixed models ((g)LMM) with the lme4 package [78] to analyze the relationship between experimental conditions and participants’ scores or reconstructed signals. The ((g)LMM) models were reported with the lme4 syntax such as:

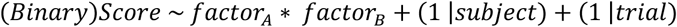

The full-factorial model is indicated by the fixed effect term 𝑓𝑎𝑐𝑡𝑜𝑟_Q_ ∗ 𝑓𝑎𝑐𝑡𝑜𝑟_R_ and includes main effects and interactions for these two main conditions. The last term of the equation describes two random intercepts: one for each subject and one for each trial, accounting for inter-individual variability and repeated measures. The significance of the fixed effect was assessed using a Chi-square test. To further investigate significant effects, post hoc pairwise comparisons were conducted using estimated marginal means (emmeans package; Lenth, 2017). Multiple comparisons were corrected using the false discovery rate (FDR) adjustment [80] to control for inflated Type I error rates.

Finally, to assess differences between the two modulation profiles, we implemented a non-parametric permutation test using a max-statistic approach to correct for multiple comparisons across scale-rate pairs, following a previously described method [81]. For the within-task analysis of MRF, the data were inherently unbalanced due to participant-dependent keyword selection. To address this, we employed a linear mixed-effects model (LMM), which effectively accommodates unequal trial numbers across responses. Multiple comparisons were corrected using Threshold-Free Cluster Enhancement (TFCE), as introduced in in earlier work [82] and extended in subsequent work [83]. Before comparing the modulation profiles between tasks, we accounted for baseline differences in global cortical MRF magnitude. Specifically, we first verified that neural tracking strength was positively associated with global cortical MRF magnitude using a linear mixed-effects model. To ensure that baseline shifts in global magnitude did not bias spectrotemporal comparisons between tasks, we then applied a mean- centering procedure across subjects and trials prior to statistical testing. This normalization ensured that observed differences reflected task-specific spectrotemporal patterns rather than baseline shifts in overall neural tracking strength. This combined LMM–TFCE pipeline yielded a statistically grounded and interpretable map of response-specific modulations, even in the presence of unequal data sampling.

## Conclusion

In conclusion, our findings demonstrate that selective attention modulates cortical speech encoding in a task-specific manner, selectively enhancing spectro-temporal features that are most relevant to the cognitive demands at hand. Rather than applying a uniform gain to the target class (in this case speech), attention acts as a flexible and adaptive filter that highlights different acoustic dimensions—such as formants during comprehension or pitch during detection—depending on the listener’s goals. This supports a discriminative model of attention, in which neural resources are dynamically allocated to serve specific perceptual goals.

A key strength of our study lies in the development of new methodological tools to study these dynamics. The introduction of the cortical Modulation Response Function (MRF) from EEG-based spectrogram reconstructions represents a significant advance in linking low-resolution neural data to interpretable sensory encoding. Despite the spatial limitations inherent to EEG, the MRF proved sufficiently sensitive to capture how attention reshapes the spectro-temporal structure of speech representations, opening a new path for studying auditory cognition under ecologically valid conditions.

Beyond speech, our findings contribute to a comprehensive insight of attention as a dynamic mechanism that align perception with cognitive goals. In modern computational models, objective functions can be explicitly defined to optimize specific features. Our data suggest that the brain operates under a similar logic: it dynamically sets internal objective functions, not at the level of sensory domains (e.g., "speech"), but within them—prioritizing the acoustic features most relevant for the current task. This perspective offers a conceptual bridge between neurophysiological data and theories of cognitive control, and can light the path for future efforts to model perception as an adaptive, goal-driven process.

## AKNOWLEDGEMENT

The authors thank Mary Shin for her assistance with data collection and stimuli creation.

## Supporting information

### Formants and pitch extraction

We extracted the acoustic modulation power spectrum (MPS) for a male and a female voice, using approximately 31 and 27 hours of audiobooks, respectively. These recordings come from three different genres (adventure, science fiction, and politics) to ensure an appropriate match in terms of prosody and other acoustic characteristics. Table S1 provides further details on these audiobooks. For each MPS, we averaged MPS calculated over 10-second segments. As illustrated in **Fig S1Error! Reference source not found.**, the MPS of the male voice is more broadly distributed across the spectral scale, whereas the MPS of the female voice is more concentrated around a modulation of 1 cycle/octave. As previously noted [32], analyzing the MPS of female voices in cycles/octave can be challenging. Indeed, the harmonic structure of female vocalic sounds creates a repeated pattern of spectral modulations. As a result, in female voices, formants and pitch do not distribute into distinct regions but rather overlap within the same spectral area. However, we know that formants primarily reside in low spectral modulations, whereas pitch is found in high spectral modulations. Thus, the observed difference between male and female MPS allowed us to identify specific areas which, in the case of male voices, correspond to the strict regions of formants and pitch.

**Table S1.** Characteristics of the six audiobooks.

| Voice | Audiobook | Duration |
| --- | --- | --- |
| Male voice (Scott Brick) | Jurassic Park [61] | 15h10min |
|  | Foundation[63] | 8h37min |
|  | Prisoners of Geography [62] | 8h49min |
| Female voice (Julia Whelan) | Congo [84] | 10h16min |
|  | The Ninth Metal [85] | 10h15min |
|  | Political Tribes [86] | 7h3min |

**Fig S1.**
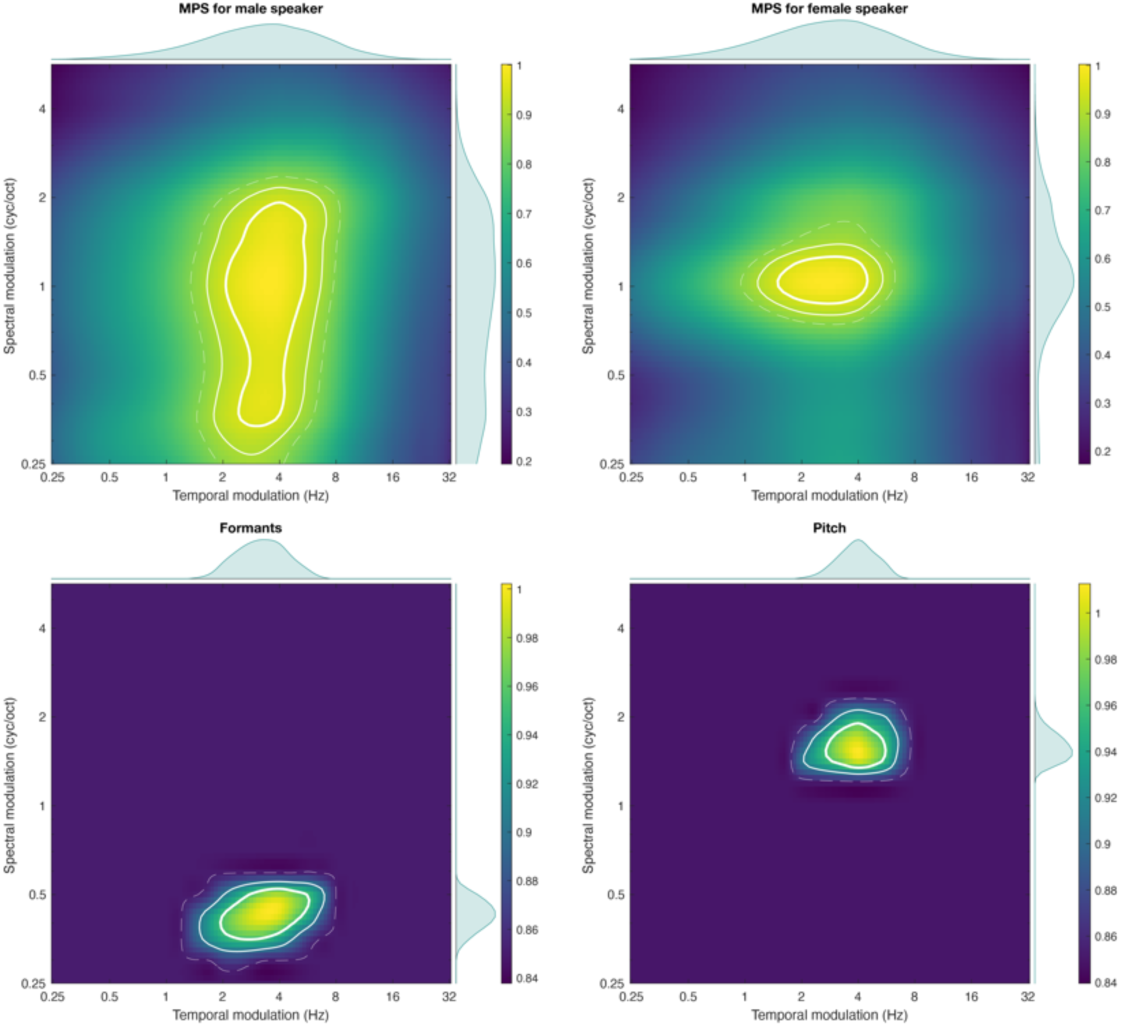
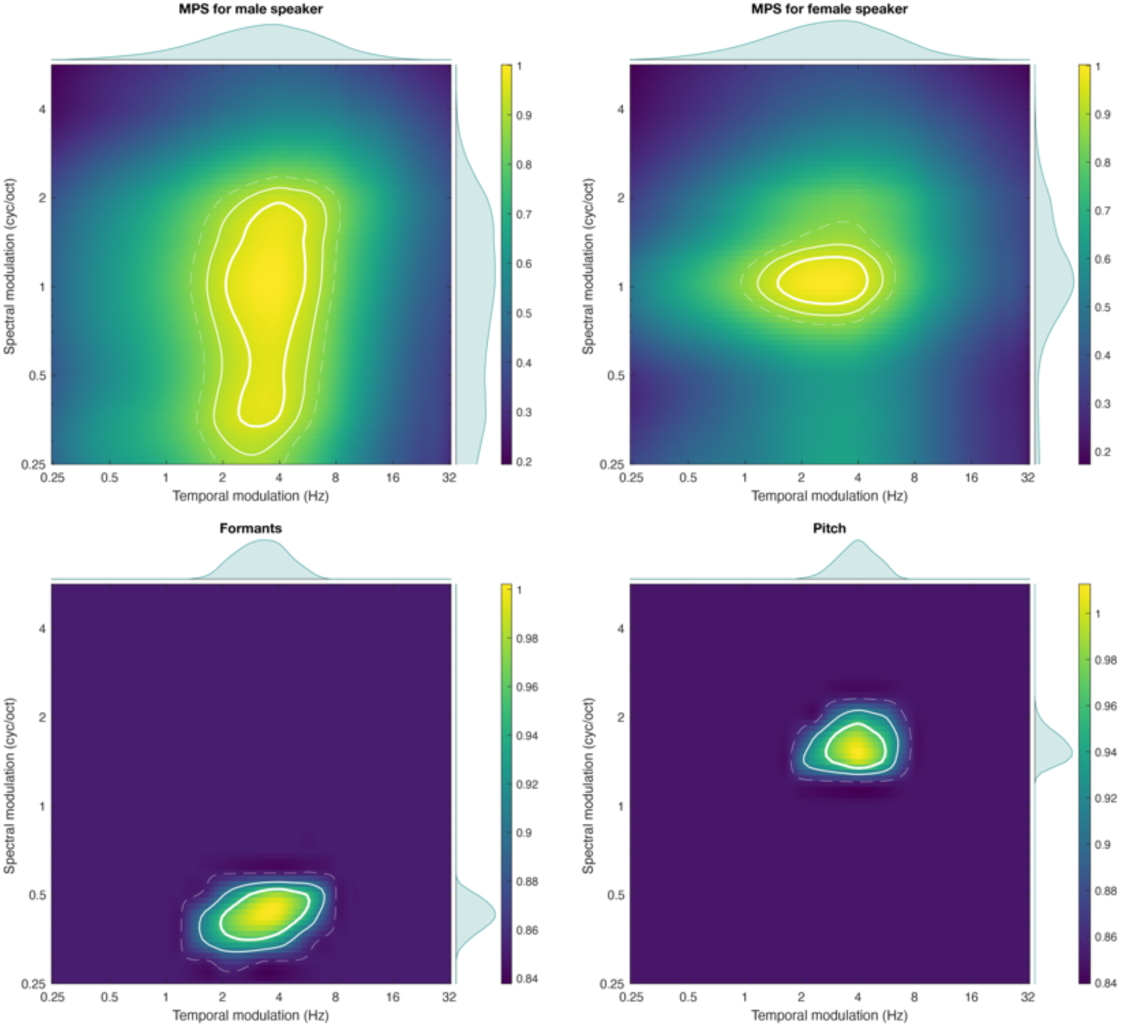
Modulation Power Spectrum (MPS) of male and female voices and distribution of formants and pitch. The figure shows the MPS computed for the male voice in the top left and for the female voice in the top right. The bottom left panel represents the distribution of formants, highlighting their presence in low spectral modulations, while the bottom right panel illustrates the concentration of pitch in high spectral modulations.

### Noise creation and validation

**Fig S2.**
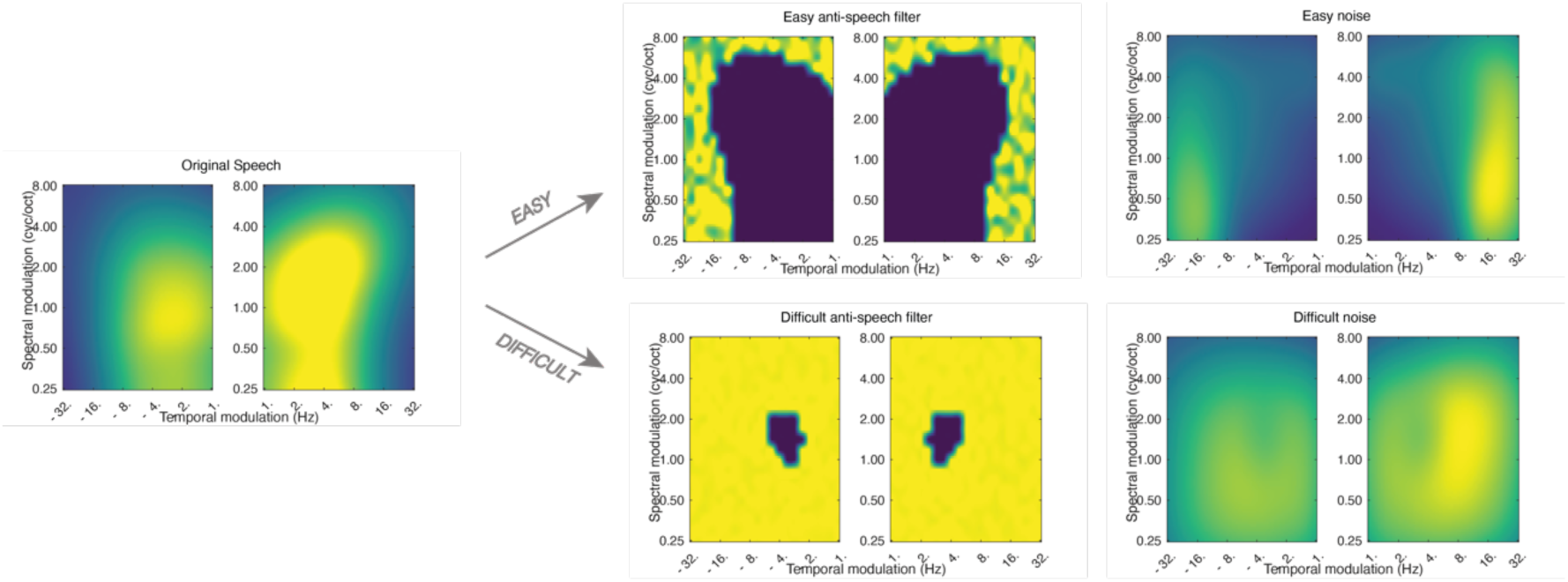
Creation of the noise. Using the NSL Toolbox, the general form of the speech was analyzed. Then a mask pro-speech was created to represent the ideal representation of the noise. Here the easy noise covers less speech than the difficult noise. The mask was then applied to a white noise cortical representation, and the noise was reconstructed accordingly.

**Fig S3.**
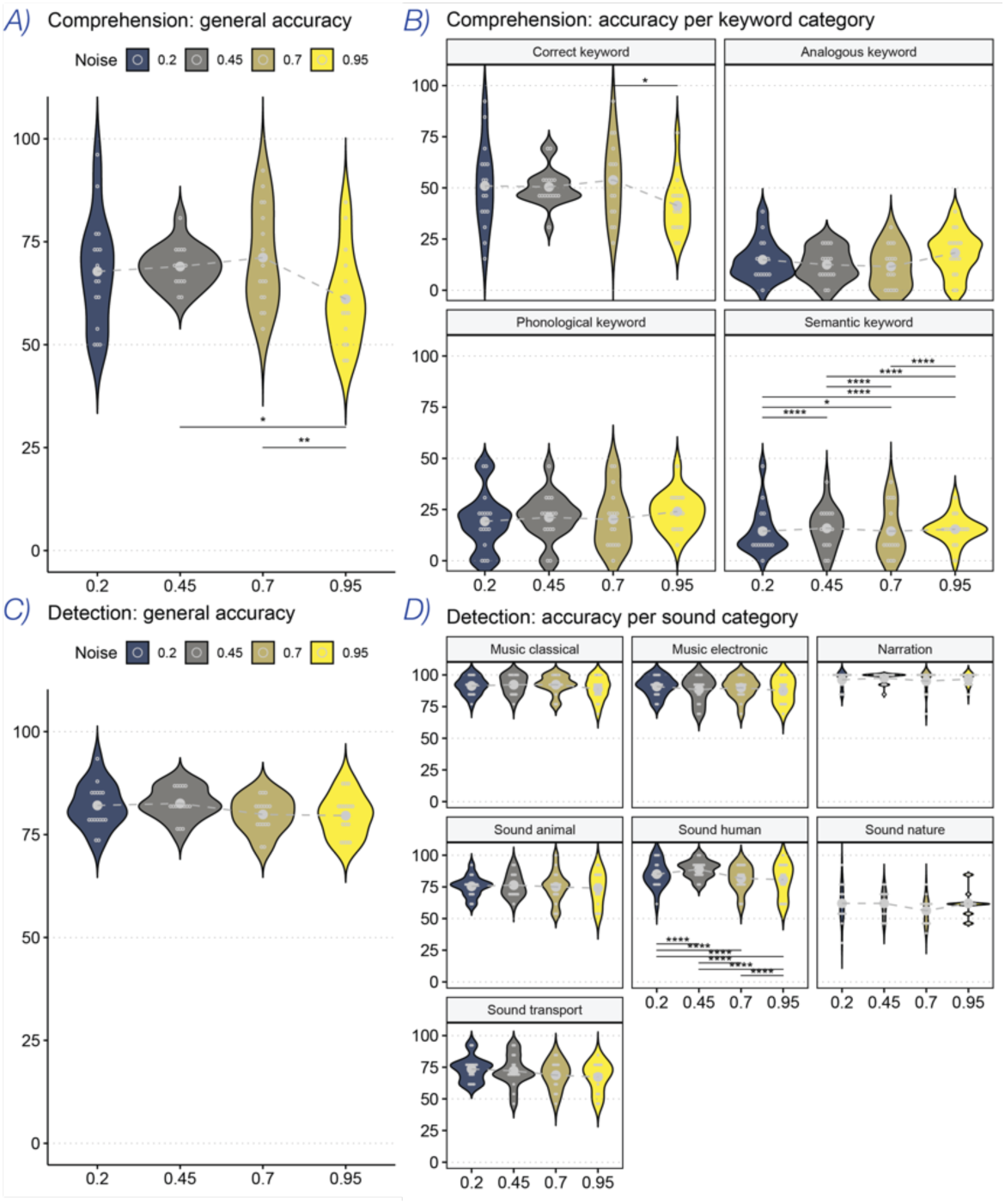
Behavioral accuracy for both tasks for preliminary studies to select the noises profiles. Panels A and B show the results for the comprehension task: A displays the overall accuracy, and B details the accuracy by keyword type. Panels C and D present the detection task: C shows overall accuracy, and D breaks down results by sound type. Theoretical chance level was 50%, with a 95% confidence interval ranging from 36% to 63%, based on a binomial distribution with 52 trials. Noise level 0.45 was the only level consistently above the chance interval for all participants for the comprehension task and was thus selected as the easier noise difficulty level. It also showed the highest detection performance, particularly driven by strong scores for human sounds. Conversely, noise level 0.95 was chosen as the difficult condition, as it showed the most significant drop in overall comprehension accuracy.

### Keywords selection

The identification of keywords in the narratives was conducted through an automated procedure that combines two dimensions of analysis: phonological and semantic. The objective was to select words with specific phonological and/or semantic relationships and to identify analogies between these categories, allowing for an assessment of cross-influences between these linguistic dimensions.

#### 1. Selection of initial keywords

Only words from the selected speech narration belonging to relevant grammatical categories (nouns, verbs, adjectives) were retained. Morphosyntactic tagging was performed using the NLTK universal tagger [87]. Words were then reduced to their canonical form using lemmatization and stemming. Finally, a lexical frequency criterion was applied to eliminate words that were too rare or too frequent in English. Frequencies were extracted from the wordfreq database [88], and only words with occurrences between 1 and 750 per million were retained.

#### 2. Selection of phonological keywords

Then, the initial keywords were compared against the CLEARPOND database [89], which provides detailed phonological information for each word. For each initial keyword, a list of phonologically similar words was generated based on the phonetic representation of each word in CLEARPOND and the calculation of phonological distance between the keyword and its phonological neighbors

#### 3. Selection of semantic keywords

The goal of this step was to identify words that share a semantic proximity with the initial keywords. To achieve this, we used the GloVe-wiki-gigaword-100 model [90], which represents words as vectors in a multidimensional semantic space. Each initial keyword was compared to other words in the corpus by computing their cosine similarity. Only words with a similarity score above 0.65 (indicating strong semantic distance) were retained for inclusion in the semantic keywords list. Additionally, phonological keywords were also evaluated in the GloVe model to exclude those that shared excessive semantic similarity with the initial keywords.

#### 4. Selection of analogical keywords

Analogical words were identified by combining phonological and semantic relationships derived in previous steps. The goal was to select words that simultaneously exhibit a phonological relationship with semantic keywords and a semantic relationship with phonological keywords. Only words with strong similarity to this newly generated representation were retained, forming the analogical keywords list. A final verification step ensured that the phonological distance between analogical keywords and phonological keywords was sufficiently large to avoid simple phonetic substitutions and the semantic distance between analogical keywords and semantic keywords was greater than their distance to phonological keywords, ensuring a true combination of both dimensions.

#### 5. Relationships between keywords

At the end of the process, four distinct keyword categories were obtained:

- Initial keywords – words extracted from the corpus, selected based on their lexical frequency and morphosyntactic relevance.
- Phonological keywords – words phonologically similar to the initial keywords.
- Semantic keywords – words semantically related to the initial keywords.
- Analogical keywords – words that simultaneously exhibit a phonological relationship with semantic keywords and a semantic relationship with phonological keywords.

### Spectrogram reconstruction

**Fig S4.**
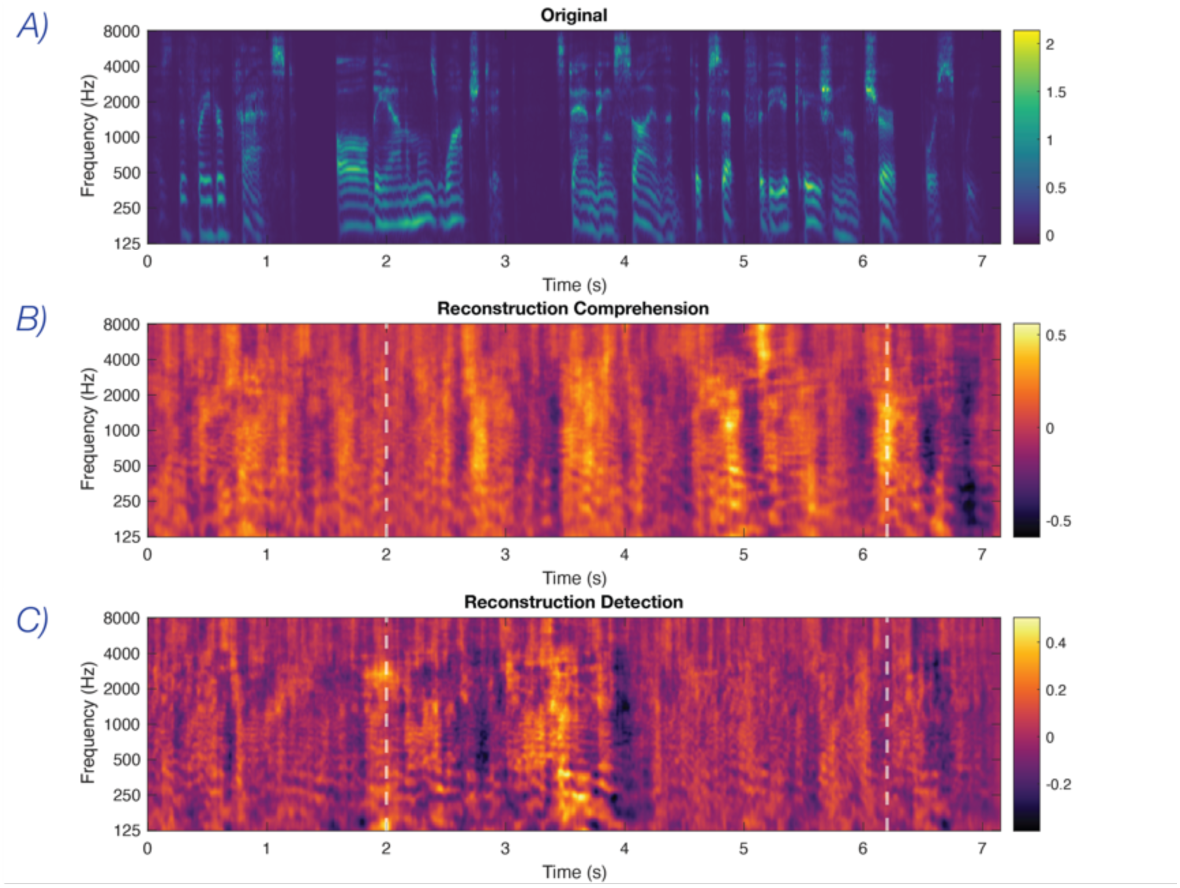
Spectrogram reconstruction and modulation profile differences across tasks. Original spectrograms (Panel A; viridis colormap) and an example of a reconstructed spectrogram for both tasks (Panels B and C; inferno colormap). Reconstructed spectrograms of the same trial using decoders trained on either the comprehension or detection task reveal distinct patterns. An example of more accurately reconstructed formants in the comprehension task can be seen around 6.15s, while an example of better-preserved harmonics in the detection task is visible around 2s. Each modulation profile is represented with temporal modulation (in Hz) on the x-axis and spectral modulation (in cycles per octave) on the y-axis.

### Head map over time

**Fig S5.**
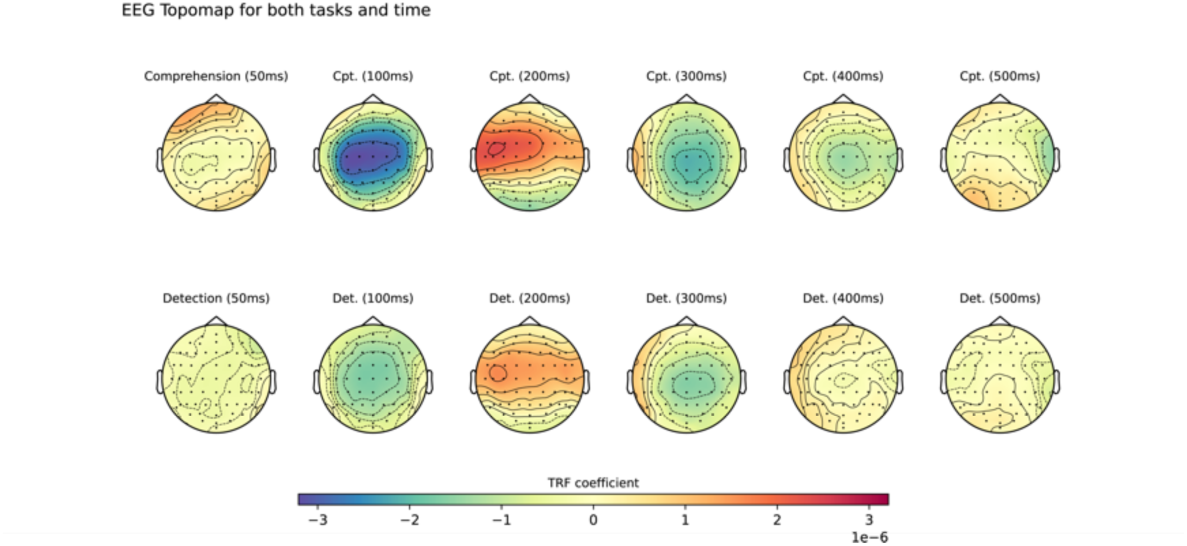
EEG topography across time lags for both tasks.

**Fig S6.**
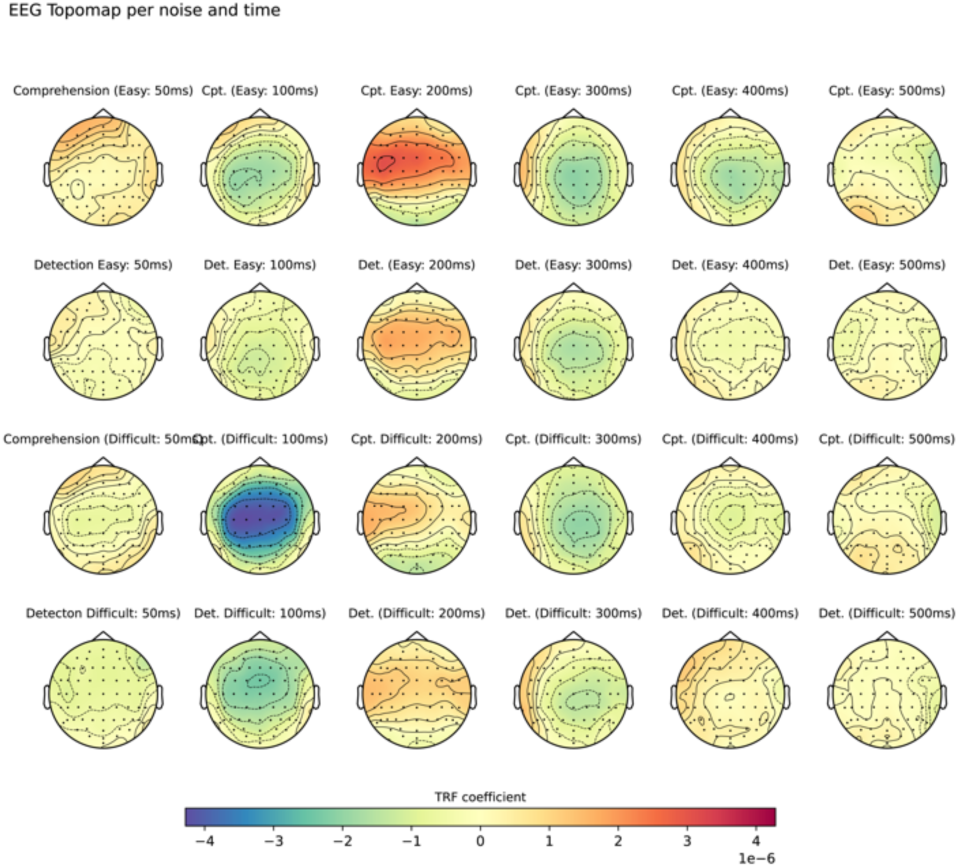
EEG topography across time lags for both tasks for each difficulty condition.

